# Transcriptional Dysregulation in the Hippocampus of a murine model for Parkinson’s Disease Cognition Impairment is Driven by Sex, Age, and Alpha-synuclein overexpression

**DOI:** 10.1101/2025.07.18.665564

**Authors:** Alessia Sciortino, Thomas Hentrich, Sergio Helgueta, Pierre Garcia, Kristopher J. Schmit, Mélanie H. Thomas, Rashi Halder, Djalil Coowar, Michel Mittelbronn, Lasse Sinkkonen, Julia Schulze-Hentrich, Manuel Buttini

## Abstract

Cognitive impairment is the most common and detrimental, but understudied non-motor symptom of Parkinson’s disease (PD). Neuropathologically, it is associated with alpha-synuclein (αSyn) misfolding and synapse loss in hippocampus and prefrontal cortex, leading to cognition loss and ultimately dementia. The molecular underpinnings of PD-associated cognitive dysfunction are unknown. In the present study, longitudinal gene expression profiling was performed to characterise molecular hippocampal alterations in a transgenic mouse overexpressing E46K mutated αSyn, a model of early PD with loss of synaptophysin, a proxy marker of cognition, in hippocampus and cortex. Comparing 4 different ages of mice from both sexes showed that hippocampal gene expression changes were sexually dimorphic and strongly modulated by age and αSyn overexpression. Pathways that emerged across different comparisons were connected to a variety of neuronal functions, collagen synthesis/remodelling, cellular stress, and inflammatory responses. The findings indicate that sex and age are essential covariates to consider when studying PD-associated cognitive decline. The uncovering of early events leading to disease in an animal model is an essential step toward prognostic biomarker identification and early interventions, which may have implications for monitoring, and for timing of therapeutic approaches.

## Introduction

Cognitive impairment is caused by a gradual reduction in memory, attention, reasoning, and language skills beyond normal aging, culminating in dementia. Dementia is characterized by a loss of cognitive function so severe that it interferes with daily functioning. Age and sex are key factors modulating brain changes not only in healthy people ^1^, but particularly in those with cognitive decline and different types of dementia ^2,3^. Age is a distinct risk factor for dementias ^3^, and sex differences have been described for most ^2^. For instance, it is well established that, in Alzheimer‘s disease dementia ^4–6^, there are differences in incidence (higher in females), age of onset (tends to occur later in females), symptoms (tends to progress faster in females), risk factors (*ApoE4* genotype-associated risk increase larger in females), and response to treatments (females tend to respond better). Studies on sex differences in other forms of dementia are sparse. Evidence suggests they exist for frontotemporal dementia ^7,8^, and for vascular dementia ^9^.

Cognitive impairment, leading to Parkinson’s disease dementia (PDD), affects around 60% of patients with Parkinson’s disease (PD) ^10,11^. PD affects over 10 million people worldwide. Non-motor symptoms, such as depression, sleep disorders, and cognitive dysfunction can manifest up to two decades before the emergence of the characteristic motor symptoms ^12,13^. A disease entity closely related to PDD is Dementia with Lewy Bodies (DLB), even though the two present some differences ^14^. Cognitive dysfunction in PD starts with mild cognitive impairment (MCI), which is present in 30% of newly diagnosed patients ^15,16^, whereas up to 50% of patients with normal cognition at baseline develop MCI after a five-year disease course ^17,18^. MCI is a strong predictor of transition to dementia ^19^. Cognitive dysfunction in PD has been associated with deficits of different neurotransmitter circuits ^10,20^. Reduced synaptic density and loss of pre-and post-synaptic markers have been shown to correlate with the extent of cognitive dysfunction in PD ^21,22^. Few studies have investigated sex differences in PD-associated cognitive dysfunction. Males were reported to have more cortical thinning ^23,24^ and some of their cognitive functions were more severely affected than in females ^23^. These observations stress the need to include age and sex as a variable in studies on cognitive decline in PD, including preclinical ones, which hasn’t been done so far.

A key tool to understand initiation and progression of neurological disease in a preclinical setting are animal models. Several rodent models have been used to investigate PD-associated cognitive dysfunction and study the hippocampus, a major brain region orchestrating cognition and memory. Spatial learning and memory retention deficits have been reported following administration of neurotoxins, as well as in αSyn (*SNCA*) overexpressing mouse models ^25,26^. One study identified 96 differentially expressed genes (DEGs) in the hippocampus of MPTP-injected mice ^27^. Gene ontology (GO) term enrichment analysis predicted impairments of DNA methylation and protein serine/threonine kinase activity. Another study investigated differential gene expression in 6-OHDA-injected rats, uncovering 45 DEGs in the hippocampus but no gene ontology (GO) terms or KEGG pathways ^28^. A recent effort investigated the spatial transcriptomic of the hippocampus of MPTP-injected mice by single-cell RNA-sequencing ^29^. The results revealed that neuronal gene expression in the hippocampus is spatially heterogeneous, and that memory deficits correlated with differential expression of synapse and calcium signalling genes in the CA1 and CA3 hippocampal subfields. Nevertheless, these studies only used male mice and one age-group, thus presenting limitations in the understanding of how PD-associated cognitive impairment evolves with aging and is influenced by sex. Hentrich and colleagues investigated hippocampal transcriptomic profiles in 6-and 12-month-old αSyn overexpressing male mice ^30^. Their results suggest that *SNCA* overexpression causes age-dependent disturbances in midlife hippocampal transcriptome, reflective of accelerated aging and a disruption of adaptation to the aging process.

To investigate the effects of sex, age, and αSyn overexpression on hippocampal transcriptional dysregulation, a longitudinal study of the hippocampal transcriptome in a mouse model overexpressing the human E46K-mutated αSyn gene under the transcriptional control of its own promoter, the BAC-Tg3(SNCA*E46K) mice ^31^, was undertaken. Four different ages were analysed, and both male and female mice, as well as their wild-type littermate controls, were used. The model was chosen for two reasons: first, the E46K mutation, is linked to the presence of early cognitive impairment ^32^, thus indicating a role of this mutation in cognitive decline; second, a previous study demonstrated the presence of several typical early PD pathologies, such as age-dependent loss of striatal tyrosine-hydroxylase and dopamine transporter, in this model ^31^, thus providing a base translational relevance to build on.

Heterozygous BAC-Tg3(SNCA*E46K) mice of both sexes developed a significant loss of synaptophysin-positive presynaptic terminals in the hippocampus and cortex, a proxy measure of cognition ^33,34^, by 14 months of age. Hippocampal gene expression profile was modulated by sex, age, and *SNCA* overexpression. Neuronal dysfunctions, immune dysregulation, vascular processes, and collagen synthesis disruption emerged as key pathways involved in aging and disease. These results indicate that PD-associated cognitive impairment is modulated by the combined effects of a multiplicity of factors that include sex, age, and abnormal αSyn. This observation may have important implications for prognostic biomarker identification and therapeutic management of this disease in a precision medicine context.

## Results

### Alpha-synuclein overexpression and synaptophysin loss in the hippocampus of BAC-Tg3(SNCA*E46K) mice

The BAC-Tg3(SNCA*E46K) mouse model overexpresses human E46K mutated αSyn in brain regions of interest for PD, including striatum, substantia nigra, hippocampus and cortex ^31^. Hippocampal human (*SNCA*) and murine αSyn gene (*Snca*) expression and human and total αSyn (SNCA) protein levels were quantified, across 4 different age groups (3, 6, 8, and 14 months), by RT-qPCR and Western Blot, respectively. *Snca* mRNA expression levels did not differ between WT and BAC-Tg3(SNCA*E46K) mice, nor between sexes, and were stable across age (Fig. 1A left). *SNCA* was expressed only in BAC-Tg3(SNCA*E46K) mice, and the expression levels were stable across ages and sexes (Fig 1A right). Similarly, human αSyn protein was detected in BAC-Tg3(SNCA*E46K) mice only (Fig. 1B right, Suppl. Fig. 1). Levels of total αSyn protein were quantified using a pan-αSyn antibody. They were stable across ages and similar in both sexes in WT mice. On average, BAC-Tg3(SNCA*E46K) mice, whether male or female, had a 4.37-fold increase (p = 2.2e-16) in total αSyn protein compared to WT mice (Fig. 1B left, Suppl. Fig. 2), indicating that the observed increase is due to the transgenic αSyn protein, and those levels did not change with age. Representative images for human and total αSyn immunofluorescence staining at all ages are shown in Fig. 1C.

**Fig. 1.**
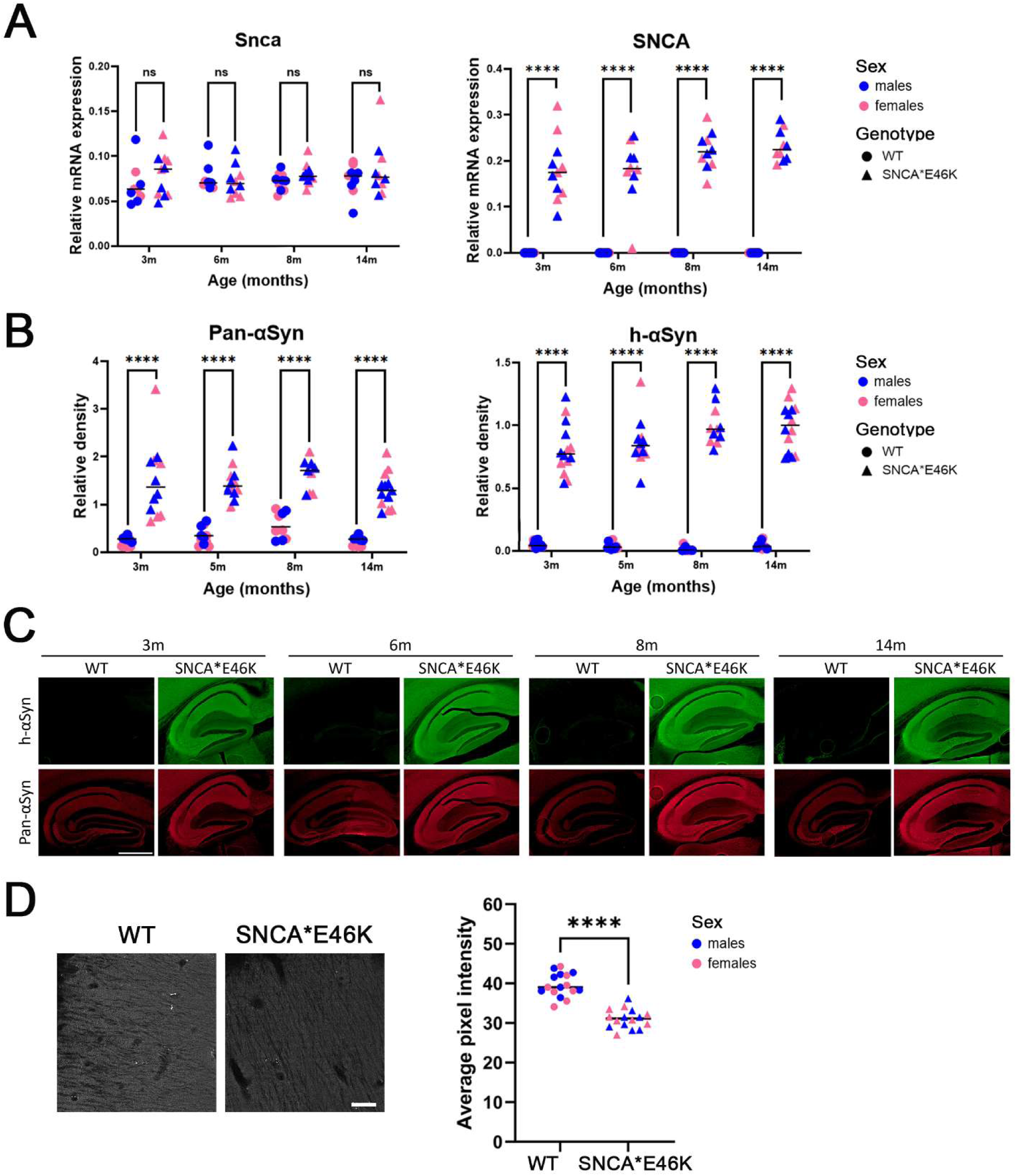
Hippocampal α-synuclein expression levels and transgene-linked synaptic degeneration in BAC-Tg3(SNCA*E46K) compared to wildtype littermates. **A)** Relative mRNA expression levels of murine αSyn (*Snca*, left) and human transgenic αSyn (*SNCA*, right) in BAC-Tg3(SNCA*E46K) (SNCA*E46K), mice and wild-type littermates (WT). No differences were observed between sexes, so both sexes were grouped for comparisons between the genotypes. As the mice aged, level of *Snca* did not change, nor did the level of *SNCA* in BAC-Tg3(SNCA*E46K) mice. No *SNCA* was detected in WT mice. N = 4 to 6 mice/group, ns = non-significant, **** = p<0.0001 (Kruskall-Wallis). **B)** Relative density of total (Pan-αSyn, left) and human (h-αSyn, right) αSyn protein in BAC-Tg3(SNCA*E46K) and WT mice. Relative density was calculated from western blot images, using β-Actin as loading control. BAC-Tg3(SNCA*E46K) mice showed a 4.37-fold increase in total αSyn levels compared to WT mice. N = 4 to 6 mice/group, **** = p < 0.0001 (Kruskal-Wallis). **C)** Fluorescent immunostaining of human (h-αSyn, green) and total (Pan-αSyn, red) αSyn protein in the hippocampus of BAC-Tg3(SNCA*E46K) and WT mice. Scale bar: 500μm. **D)** Left panel: Representative images for Synaptophysin (SYP) immunostaining in the hippocampal pyramidal layer of 14-month-old BAC-Tg3(SNCA*E46K) and WT mice (Scale bar: 50 μm). Right panel: SYP staining intensity quantification in the hippocampal pyramidal layer of 14-month-old BAC-Tg3(SNCA*E46K) and WT mice. N = 8 mice/group, **** = p < 0.0001 (Student’s t).

Synaptophysin, a synaptic vesicle protein whose loss correlates with cognitive impairment in Alzheimer’s disease ^35^, and has been shown to reflect true synapse loss in a model thereof ^36^, was quantified in hippocampus and cortex. At 14 months of age, both male and female BAC-Tg3(SNCA*E46K) mice showed a significant and similar loss of synaptophysin in the pyramidal layer of the hippocampus and in the cortex (layer 2-5) compared to their WT littermates (Fig. 1D, Suppl. Fig. 3). These results show that transgenic αSyn overexpression is sufficient to induce synaptic terminal damage, a correlate of cognitive dysfunction, in 14-month-old mice. Interestingly, despite the loss of synaptophysin, no αSyn intraneuronal inclusions or aggregates were observed in the hippocampus with the αSyn antibodies used (see above). Toxic αSyn oligomers could be the cause for synaptic degeneration ^37^. Alternatively, αSyn overexpression itself, by inhibiting neurotransmitter release ^38^, could cause this degeneration ^39^. The investigation of this issue is however not the focus of the present study.

Overall, these results show that the BAC-Tg3(SNCA*E46K) mouse is a suitable model to study the molecular events in the hippocampus associated with synaptic injury.

### Quality control of sequencing data and identification of gene expression drivers

Gene expression profiling of the hippocampus of BAC-Tg3(SNCA*E46K) and WT mice was performed at 3, 6, 8, and 14 months of age by bulk RNA-sequencing (RNAseq) analysis. Bulk RNAseq is a well-established, cost-effective technique that allows robust detection of differential gene expression. The lower noise and variability of bulk versus single-cell expression data, combined with increased statistical power, make bulk RNAseq, in a first step, well-suited for capturing dynamic changes in gene expression across different conditions or time points ^40^.

To check for cell type variability in sample composition, the representation of different cellular identities across samples was investigated. Cell type-specific expression profiles were obtained from reference profiles ^41^ and plotted for all samples individually (Suppl. Fig. 4). Overall, no differences were observed in cell type-specific expression profiles, indicating that samples had homogeneous cellular composition. To identify closely related clusters within the datasets, principal component analysis (PCA) was performed on the raw data. The samples partitioned into four equidistant groups, whose separation could be attributed to sex and an unknown factor (Suppl. Fig. 5). The separation of samples along the x axis could not be attributed to experimental factors such as housing of the mice, or RNA extraction day (not shown). We assume that the hybrid strain background on which the mice were bred was responsible for the observed separation of the samples ^42^. Nevertheless, determining the molecular features of strain-specific characteristics was beyond the scope of this study. Variability due to genetic background is also typical for human study populations. Therefore, an unrestricted surrogate variable analysis was used to correct the data. After correction, the samples separated into four equidistant clusters based on sex and genotype (Fig. 2). Furthermore, within each cluster, the samples appeared to be distributed along a gradient that reflected age. Overall, the PCA analysis revealed that sex, genotype, and age congruently drove gene expression dysregulation in the hippocampus of BAC-Tg3(SNCA*E46K) mice and their WT littermates. Therefore, each of these components was taken as a separate starting point for in-depth follow-up analysis.

**Fig. 2.**
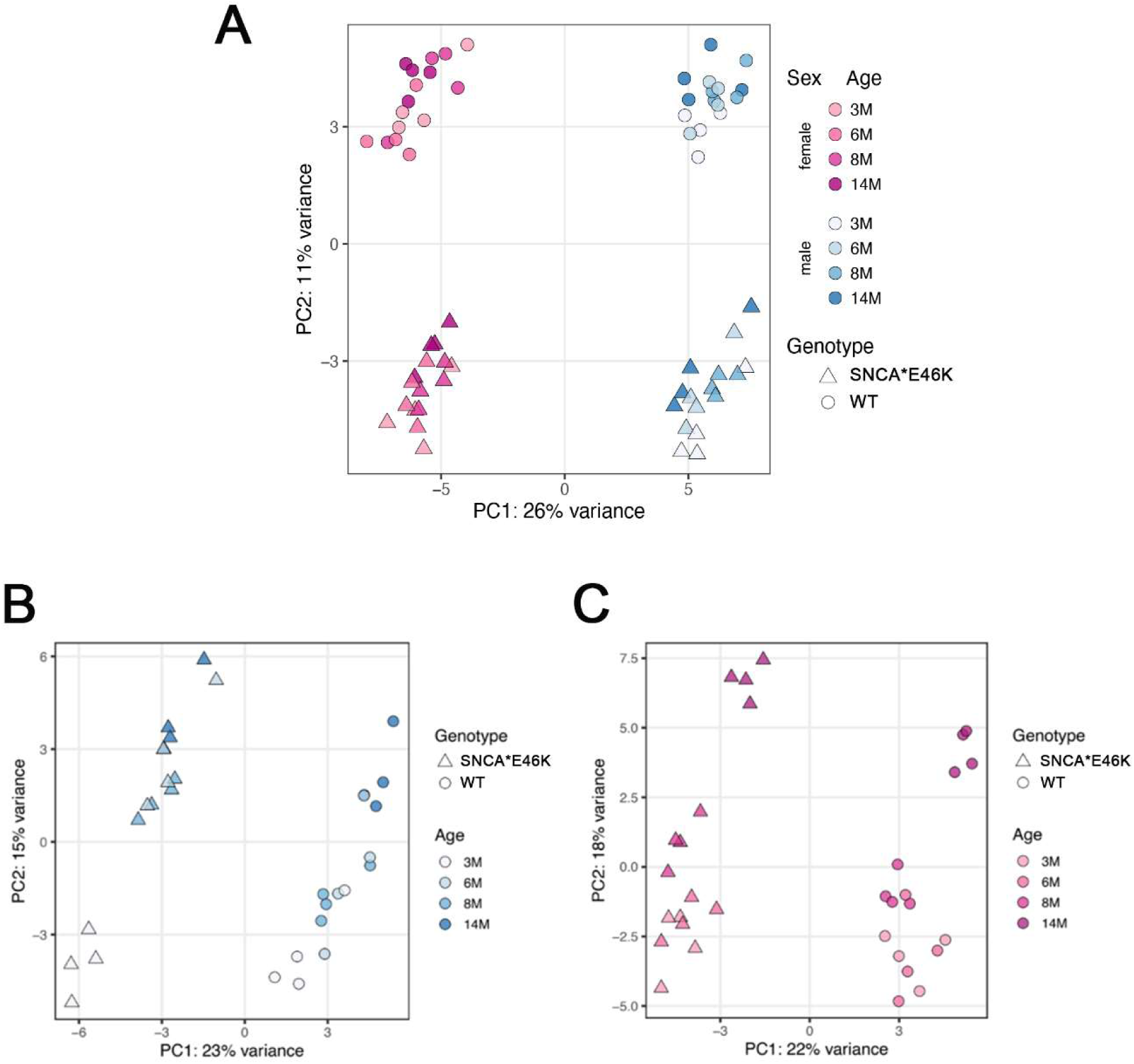
**Hippocampal gene expression profiles in BAC-Tg3(SNCA*E46K) mice and WT littermates were driven by sex, age, and genotype**. **A)** Dimensionality-reduced representation (Principal Component analysis) of hippocampal gene expression profiles for all samples after bulk RNA sequencing. First and second component of the principal component analysis on the top 500 most variable genes shown. Expression data corrected with sva (*sva*, v3.44). Samples cluster according to sex and genotype, and further distribute along an age-dependent gradient. N = 4 mice/group. **B)** Similar to A, for males only. **C)** Similar to A, for females only. Note the separations based on genotype, and the age gradient, in both sexes.

### Transcriptional dysregulation driven by sex

Sex-dependent gene expression differences were investigated separately in BAC-Tg3(SNCA*E46K) and WT mice, and, within each genotype, separately for each age group. For each comparison, the number of differentially expressed genes (DEGs) and the top 10 canonical pathways are shown in Fig. 3.

**Fig. 3.**
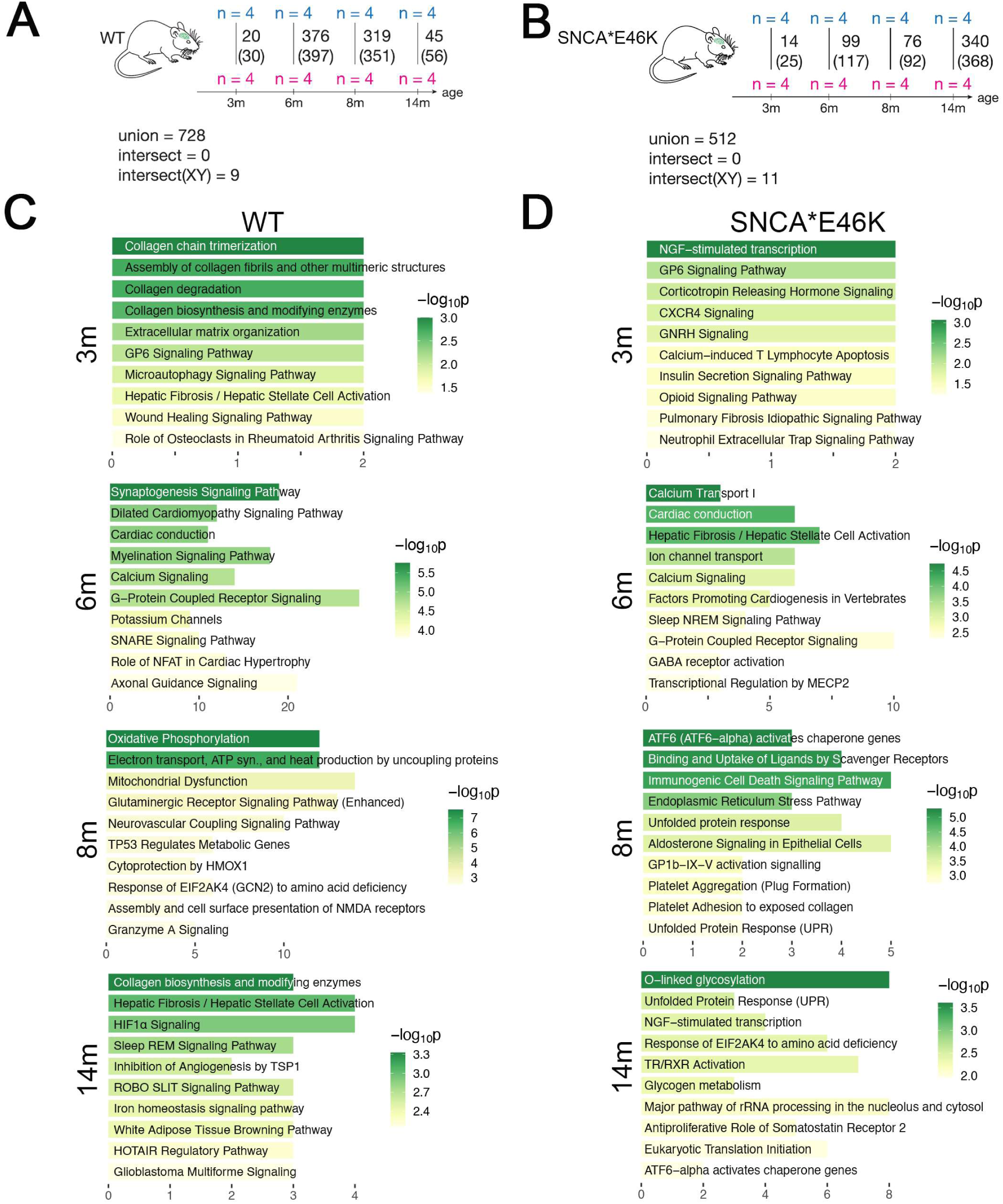
**Differentially expressed genes and pathway analysis of hippocampal gene expression profiles modulated by sex, separately in BAC-Tg3(SNCA*E46K) and WT mice**. **A)** Number of differentially expressed genes (DEGs) between male and female WT mice across age. Numbers in brackets indicate DEGs including those from sex chromosomes, numbers without brackets indicate DEGs on autosomes only. The number of all unique DEGs across all age groups (union), and the number of common DEGs across age groups (intersect) is also shown. N = 4 mice/group. **B)** Similar to A, for BAC-Tg3(SNCA*E46K) mice. **C)** Top ten IPA canonical pathways ranked by p-value in WT mice associated to sex, for each age. The number of DEGs identified in each pathway are displayed on the x-axis. **D)** Similar to C, for BAC-Tg3(SNCA*E46K) mice. See main text for details.

To control for the absence of influence of the oestrous cycle in the analysis of sex-dependent DEGs, the overall variance in male *versus* female samples was calculated with a pair-wise Pearson correlation based on the variance-stabilised counts for 43015 genes (not expressed genes were removed). Male and female samples showed a comparable distribution of sample-to-sample correlations, indirectly excluding a relevant impact of the oestrous cycle on hippocampal gene expression in female mice (Suppl. Fig. 6).

In WT mice, the sex-dependent numbers of DEGs were highest at 6 and 8 months, whereas in BAC-Tg3(SNCA*E46K) mice, this was the case at 14 months. Expression of sex-linked genes, such as *Xist* and *Kdm5d*, was clearly distinct between male and females in each genotype and at each age (Suppl. Fig. 7). Autosomal genes on the other hand showed dynamic expression patterns that changed with age in each genotype. Surprisingly, within each genotype, no autosomal DEGs were shared across the different ages (“intersect = 0”) and only a limited number of sex-linked genes (9 in WT mice, and 11 in BAC-Tg3(SNAC*E46K) mice) were common to all age groups (Fig. 3A-B).

The 11 sex-linked DEGs identified in BAC-Tg3(SNCA*E46K) mice included the 9 DEGs detected in WT mice, plus *Ddx3x* and *Kdm5c* genes. The expression pattern of these two genes in WT mice though was similar to that in BAC-Tg3(SNCA*E46K) mice, and the differences in WT mice just closely failed to reach significance.

Thus, sex-linked genes are unlikely to be essential in the ageing and disease processes.

Canonical pathways were identified for the different comparisons using the Ingenuity Pathway Analysis (IPA) software (QIAGEN). The top 10 canonical pathways for each comparison were plotted based on their-log_10_p value and the number of DEGs involved (Fig. 3). Pathway predictions based on 5 or less DEGs were excluded. There was little overlap in canonical pathways across comparisons, indicating that sex-dependent transcriptional regulation is a highly dynamic and heterogeneous process. In WT mice, synaptogenesis and calcium signalling were the pathways that differed most significantly between the sexes at 6 months, while at 8 months oxidative phosphorylation and mitochondrial dysfunction were more relevant. Calcium signalling was identified in 6-month-old BAC-Tg3(SNCA*E46K) mice, while in older transgenic mice the most significant pathway was O-linked glycosylation.

When looking at overarching functional clusters (see Materials and Methods), the most interesting observation was that BAC-Tg3(SNCA*E46K) males and females, at 8 and 14 months, differed in their stress/anti-oxidant/protein misfolding responses (Table 1), indicating differences in protective processes in males *versus* females in response to abnormal αSyn.

**Table 1:**
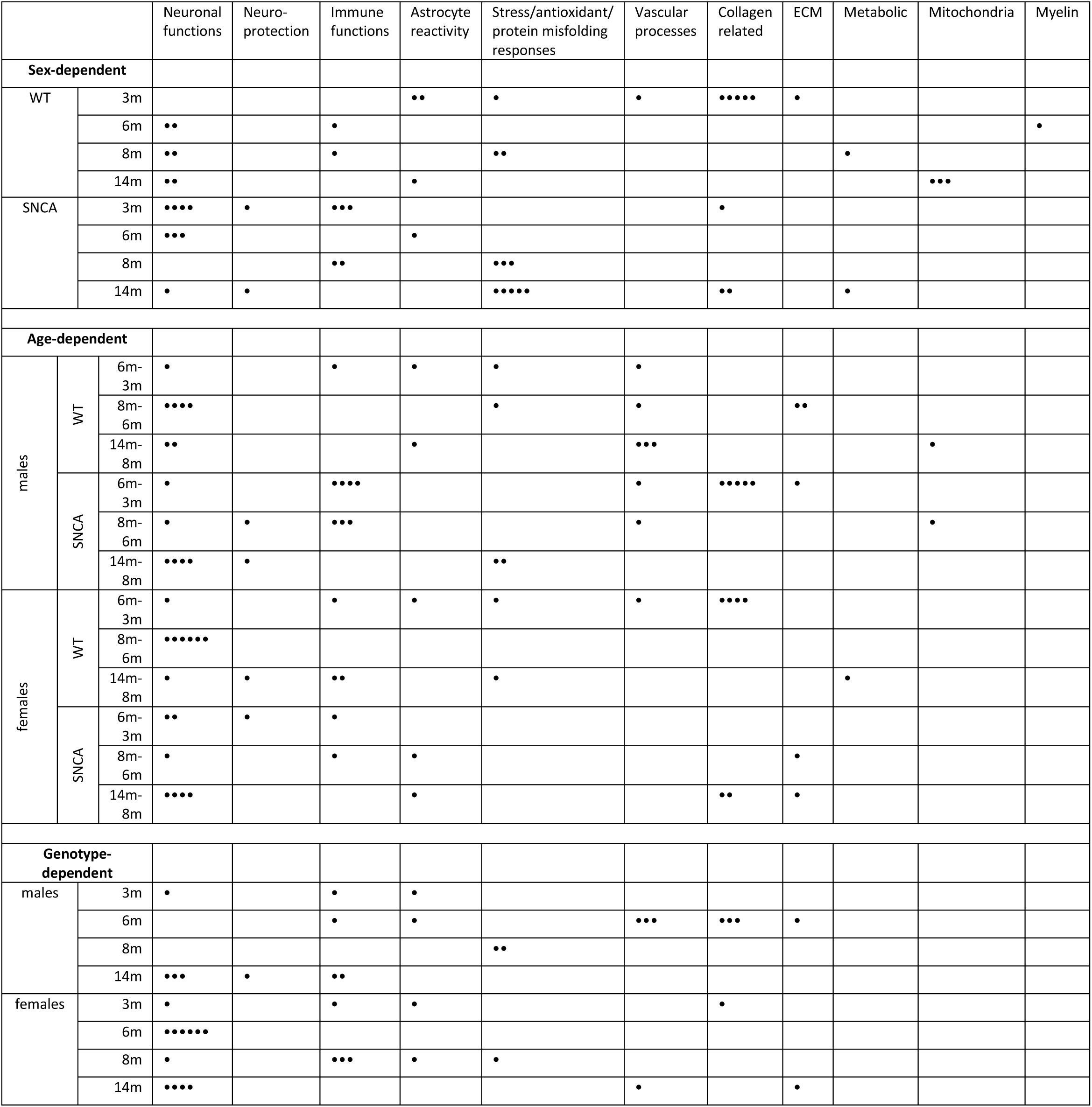
Functional clusters associated with sex-dependent (upper rows), age-dependent (middle rows) and genotype-dependent (lower rows) differences in hippocampal gene expression profiles of wild-type (WT) and BAC-Tg3(SNCA*E46K) mice. Pathways identified by IPA were manually grouped into functional clusters. For each pairwise comparison of hippocampal gene expression profiles (sex-, age-or genotype-dependent), a“●” was added to a cluster table cell each time a pathway attributed to that cluster appeared in the IPA analysis of that comparison. See main text for details.

In summary, sex-dependent gene expression differences in BAC-Tg3(SNCA*E46K) versus WT mice across different ages, revealed similar patterns for sex-linked, but different ones for autosomal genes. These differences pointed to dynamic transcriptional regulation and significant variation in specific functional clusters (e.g. stress responses) between male and female mice.

### Transcriptional dysregulation driven by age

Age is the biggest known risk factor for PD and dementia ^15,43^.

Aging is associated with shifts in brain gene expression in humans and mice ^44,45^. A recent study ^46^ highlighted a set of 82 downregulated genes in 10 or more brain regions of aging mice, which were combined into a“common ageing score” (CAS). The CAS was used here to investigate ageing patterns of gene expressions in the hippocampus of BAC-Tg3(SNCA*E46K) and WT mice (Suppl Fig. 8). A robust ageing signature was identified for all mice, despite expression patterns being noisier in males. At this level of analysis though, no clear difference was observed between WT and BAC-Tg3(SNCA*E46K) mice. This high-level view of CAS signature may not be sensitive enough to pick up gene expression changes between the two genotypes.

To better investigate the progression of gene expression changes over time, hippocampal transcriptome in 6-vs. 3-month-old, 8-vs. 6-month-old, and 14-vs. 8-month-old mice were computed for each genotype and sex separately. In males (Fig. 4), the highest number of DEGs in WT mice was observed at 6 vs. 3 months, and in BAC-Tg3(SNCA*E46K) mice at 14 vs. 8 months (Fig 4A). In females (Fig.5), the highest number of DEGs in WT mice was observed at 8 vs. 6 months, and in BAC-Tg3(SNCA*E46K) mice, similar to males of the same genotype, at 14 vs. 8 months (Fig. 5A).

**Fig. 4.**
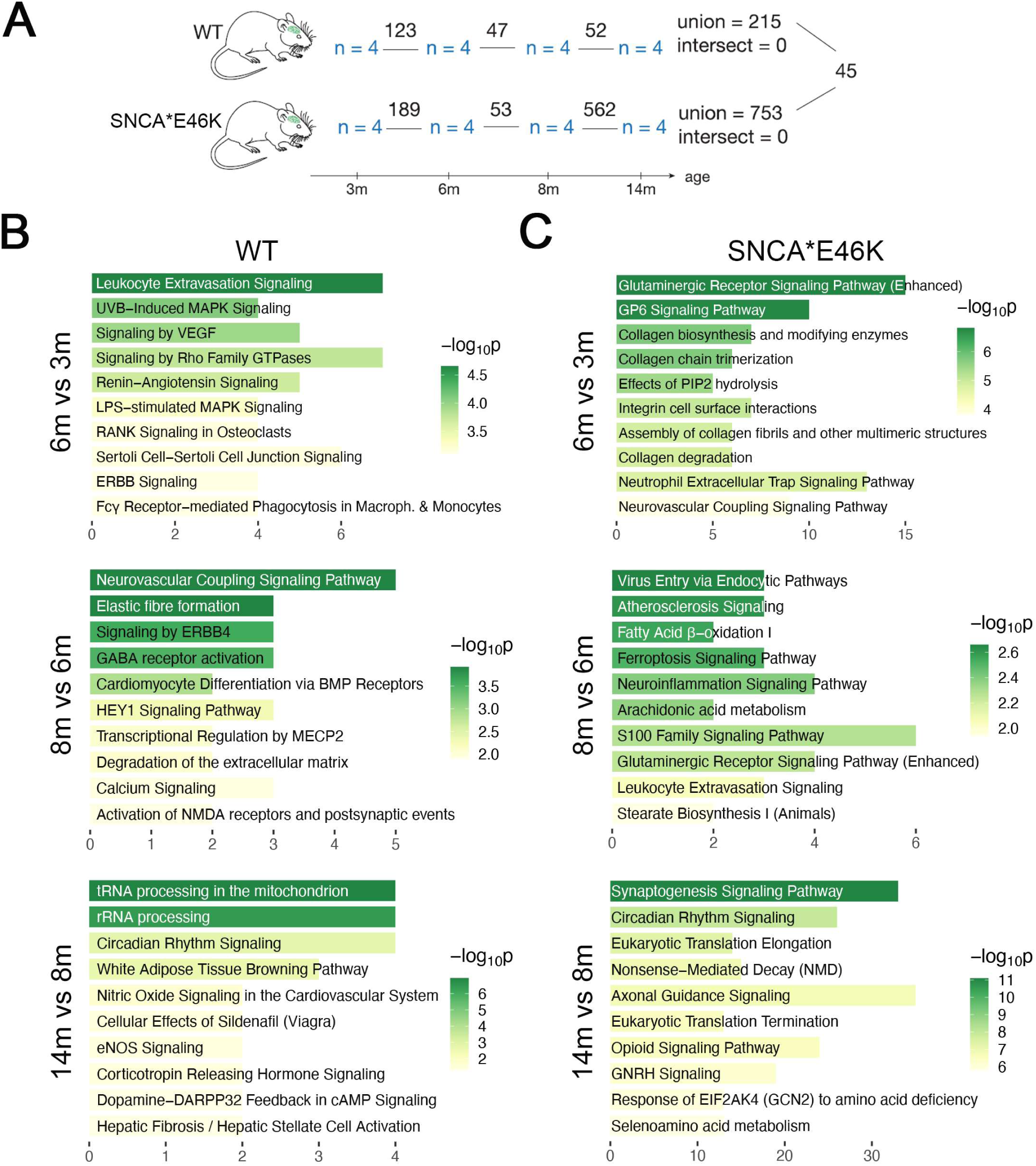
**Differentially expressed genes and pathway analysis of hippocampal gene expression profiles modulated by age in male BAC-Tg3(SNCA*E46K) and WT mice**. **A)** Number of DEGs between consecutive ages for BAC-Tg3(SNCA*E46K) and WT male mice. The number of all unique DEGs across all age groups (union), and the number of common DEGs across all age groups (intersect) is also shown. Additionally, the number of common DEGs between the two genotypes is shown on the right. **B)** Top ten IPA canonical pathways ranked by p-value in male WT mice for each comparison. The number of DEGs identified in each pathway are displayed on the x-axis. **C)** Similar to B, for BAC-Tg3(SNCA*E46K) male mice. See main text for details.

**Fig. 5.**
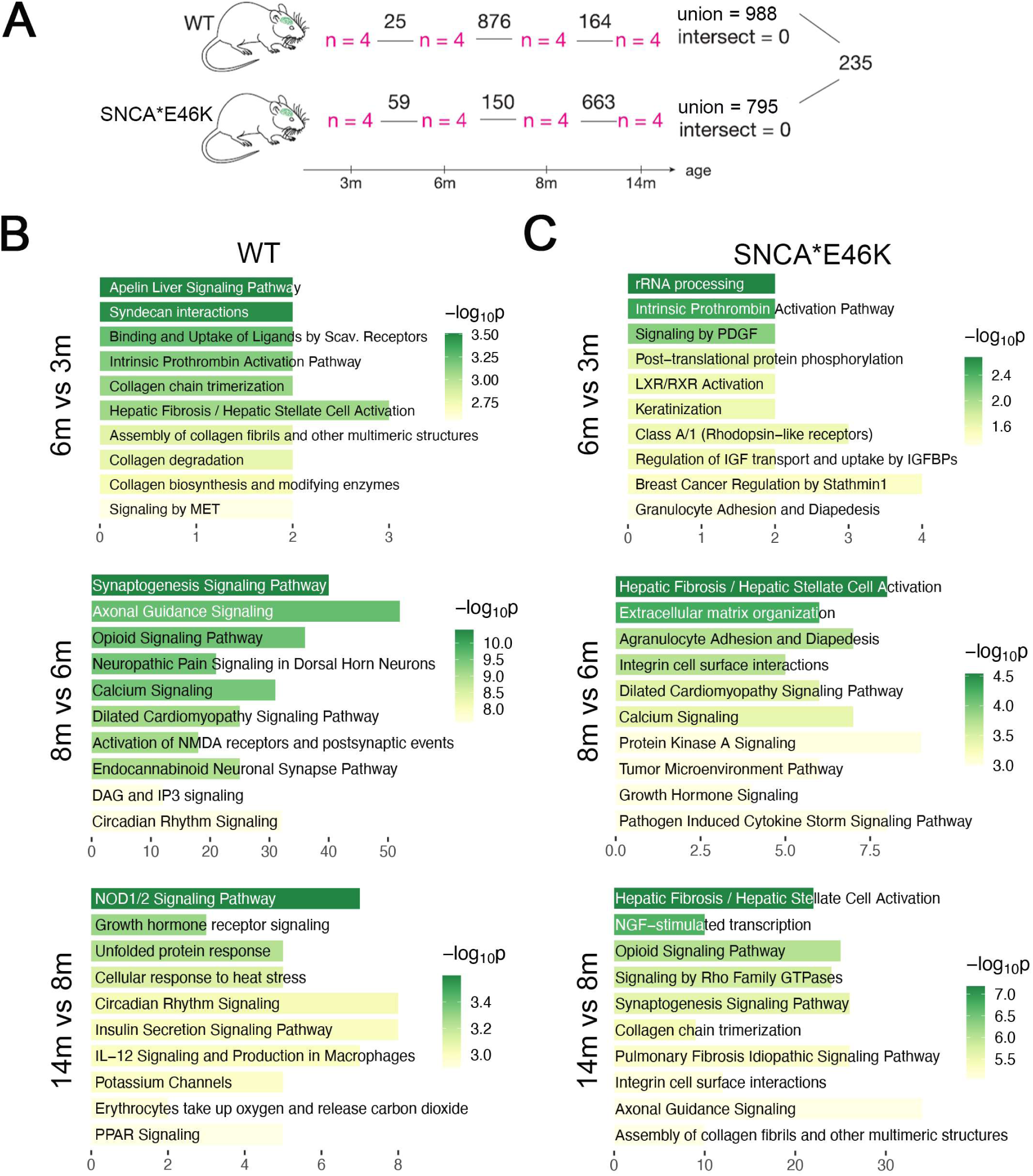
Differentially expressed genes and pathway analysis of hippocampal gene expression profiles modulated by age in female BAC-Tg3(SNCA*E46K) and WT mice. **A)** Number of DEGs between consecutive ages for BAC-Tg3(SNCA*E46K) and WT female mice. The number of all unique DEGs across all age groups (union), and the number of common DEGs across all age groups (intersect) is also shown. Additionally, the number of common DEGs between the two genotypes is shown on the right. **B)** Top ten IPA canonical pathways ranked by p-value in WT female mice for each comparison. The number of DEGs identified in each pathway are displayed on the x-axis. **C)** Similar to B, for BAC-Tg3(SNCA*E46K) female mice. See main text for details.

In total, only 215 age-dependent DEGs were identified in male WT mice across all ages (“union”), while the overall number of DEGs across all ages was over 3 times higher in all other groups: 753 DEGs in BAC-Tg3(SNCA*E46K) males, 988 DEGs in WT females, and 795 DEGs in BAC-Tg3(SNCA*E46K) females. Out of these, 45 DEGs were common between WT and BAC-Tg3(SNCA*E46K) males, while 235 DEGs were shared by females of both genotypes. For male and female mice of both genotypes, no DEGs were shared across comparisons (“intersect = 0”), indicating that age-dependent DEGs are specific to each time point, and therefore that hippocampal gene expression is highly dynamic during ageing in these mice.

Canonical pathways analysis revealed a complex picture, with some pathways being genotype-specific and others occurring in both BAC-Tg3(SNCA*E46K) and WT mice either at the same or at different ages. Synaptogenesis signalling, opioid signalling and axonal guidance pathways were changed in BAC-Tg3(SNCA*E46K) mice of both sexes at 14 vs. 8 months, as well as in WT females at 8 vs. 6 months. Significantly altered pathways in 6-vs. 3-month-old male mice included MAPK and Rho family GTPases signalling in WT, and glutaminergic receptor signalling, GP6 signalling and collagen biosynthesis in BAC-Tg3(SNCA*E46K). Calcium signalling was significant in 8 vs. 6 months female mice of both genotypes. Additionally, NMDA signalling appeared in 8 vs. 6 months WT females, while integrin cell surface interactions were identified in BAC-Tg3(SNCA*E46K) females at 8 vs. 6 months, and 14 vs. 8 months.

When looking at overarching functional clusters, the majority of the age-dependent gene expression changes occurred at the level of neuronal functions, across multiple comparisons, including, interestingly, the comparison of the oldest age groups (14 vs. 8 months) in BAC-Tg3(SNCA*E46K) mice of both sexes (Table 1). This indicates enhanced neuronal activities, notably in synapses and axons, in response to αSyn overexpression. Gene expression changes related to immune functions showed enhanced involvement in male BAC-Tg3(SNCA*E46K) mice at younger ages (6 vs. 3 months, 8 vs. 6 months).

In summary, aging causes significant shifts in brain gene expression in both WT and BAC-Tg3(SNCA*E46K) mice. Analysis of age-dependent gene expression changes revealed dynamic patterns specific to each time point and sex, with significant variations in neuronal and immune functions between the genotypes.

### Transcriptional dysregulation driven by transgenic **α**Syn overexpression

Transgenic αSyn driven gene expression changes were investigated for each sex and each age separately. The number of DEGs between the two genotypes in males at any age was comparatively low, with the highest number for a single comparison being 62 DEGs at 6 months, and an overall number of 140 DEGs (“union”, Fig. 6A). On the other hand, the comparison of the two genotypes in female mice yielded 526 DEGs overall, with the highest number of DEGs at 6 and 14 months (259 and 299, respectively, Fig. 6B). For both sexes, each comparison yielded its own set of DEGs, and no DEGs were shared across all ages (“intersect”).

**Fig. 6.**
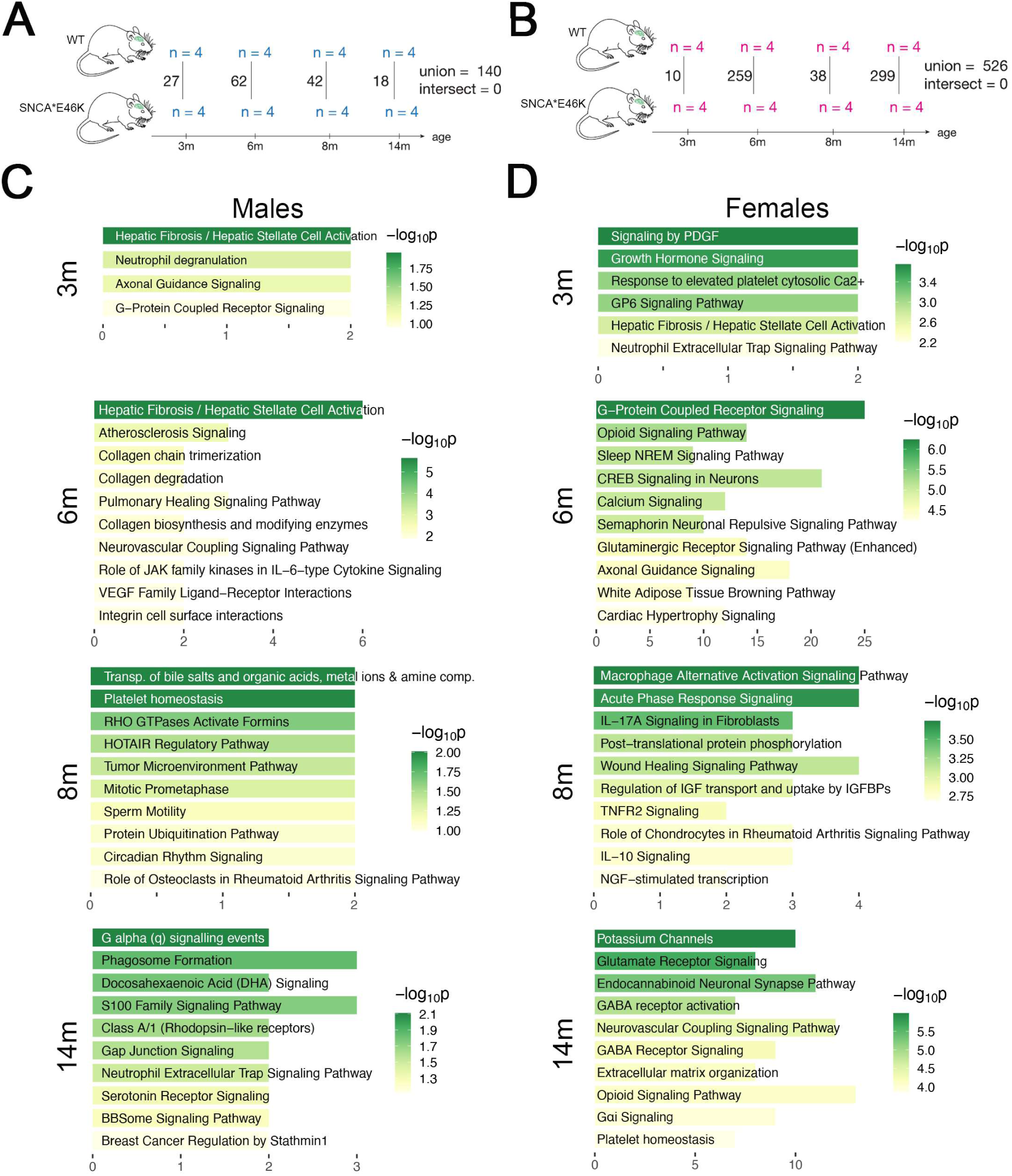
Differentially expressed genes and pathway analysis of hippocampal gene expression profiles modulated by genotype in BAC-Tg3(SNCA*E46K) *versus* WT mice. A) Number of DEGs between BAC-Tg3(SNCA*E46K) and WT male mice for each age. The number of all unique DEGs across all age groups (union), and the number of common DEGs across all age groups (intersect) is also shown. N = 4 mice/group. **B)** Similar to A, for female mice. **C)** Top ten IPA canonical pathways ranked by p-value in male mice at each time point. The number of DEGs identified in each pathway are displayed on the x-axis. **D)** Similar to C, for female mice.

Canonical pathways differences between genotypes varied across ages in males and females, but were unremarkable in male mice, in agreement with their low number of DEGs. In female BAC-Tg3(SNCA*E46K) mice, several signalling pathways were enriched at 6 months, including G-protein coupled receptor, opioids, CREB, semaphorin, glutaminergic receptor and calcium signalling. At 14 months in these mice, most affected pathways included glutamate receptor and GABA receptor signalling, as well as potassium channels.

Looking at overarching functional clusters, αSyn overexpression drove most gene expression changes related to neuronal activities in 6-and 14-month-old female BAC-Tg3(SNCA*E46K) mice, and some in 14-month-old male BAC-Tg3(SNCA*E46K) mice (Table 1). Just as seen in the gene expression changes driven by age (see above), αSyn overexpression increased genes linked to neuronal activities in the oldest age of BAC-Tg3(SNCA*E46K) mice, possibly as a driver of or response to hippocampal synaptic degeneration.

Of note are the gene expression changes related to collagen that were detected in different comparisons. In the CNS, collagen is more abundant in the basement membranes of blood vessels, providing structural support for BBB, than in the ECM ^47^. Together with the gene expression difference observed under the functional cluster of vascular processes (Table 1), this raises an intriguing possibility that sex, age, and abnormal αSyn alters vascular functions and that this may contribute to hippocampal structural or functional alterations as a consequence of aging and PD, a connection for which some evidence already exists ^48^.

In summary, the results showed that BAC-Tg3(SNCA*E46K) female mice had substantially more DEGs than their male counterparts. The gene expression changes in female mice involved pathways related to neuronal activity and vascular functions, suggesting that αSyn overexpression may contribute to hippocampal synaptic degeneration and vascular alterations associated with aging and disease in females.

## Discussion

Cognitive impairment is a key non-motor symptom of PD ^10^. Because of its significant impact on prognosis and quality of life, neurologists regularly assess it in PD patients ^49^. A closely related neurological affliction is Dementia with Lewy Bodies (DLB). While both have many overlapping features, classically in PDD, PD-typical motor symptoms precede the appearance of cognitive decline, whereas in DLB, cognitive decline occurs either before or together with the onset of motor symptoms ^50^. More recently though, arguments have been put forward that PDD and DLB exist on a continuous spectrum ^51^, and may just be different manifestations of the same disease ^52^.

Despite its socio-economic burden, PD-associated cognitive impairment is understudied compared to other forms of dementia, such as Alzheimer’s dementia, both at the preclinical and the clinical level ^53,54^. One reason for this may be a lack of awareness among the public and healthcare professionals. The lack of knowledge leads to difficulties in diagnosing it, compounded by the fact that it can be misdiagnosed for other forms of dementia.

Abnormal αSyn, neuroinflammation, neuronal dysfunctions, genetic predisposition factors, concomitant aggregation of amyloid-beta or tau, and vascular dysfunction may all contribute to the development and progression of PD-associated cognitive impairment ^54,55^.

Uncovering early molecular underpinnings of neurodegeneration is crucial for discovering prognostic biomarkers and for a better understanding of the disease. Animal models are a useful tool in that endeavour. Therefore, in this study, a model of early PD was used ^31^ to study molecular changes linked to PD-associated cognitive impairment. This model overexpresses E46K mutated αSyn in the brain, notably in the hippocampus and cortex, and this was linked to loss of presynaptic synaptophysin (see results), a proxy marker for cognition in dementias, including PDD ^33^.

Longitudinal gene expression profiling of the hippocampus in that model, and its controls, uncovered effects of sex, age, and genotype. Interestingly, the DEGs emerging from the different comparisons showed no overlap, indicating a very dynamic nature of gene expression changes in the hippocampi of these animals.

Pathways that emerged in the different comparisons indicated activities in various cellular functions, notably neuronal and immune, stress responses, and vascular activities. They also highlighted differences between sexes, across ages, and between genotypes. Because mice in this study were profiled from a young age (3 months) up to an age of over a year (14 months), the results show that molecular events in the hippocampus of this model for early PD-linked cognitive impairment differ between males and females, and highly depend on age. While further studies need to be conducted in additional PD models, the results of this study indicate that studies aimed at discovery of prognostic biomarkers of PD-associated cognitive decline need to include sex and age as covariates.

A couple of studies describe hippocampal gene expression profiles in rodent models (see Introduction), but there is only one study describing gene expression profiles of different brain regions, including hippocampus, in PD versus controls ^56^. This study also includes Alzheimer’s disease. The study found that twelve genes were altered similarly in both diseases, while four genes exhibited differential expression. Important for this discussion, the study identified several genes that were altered in the hippocampus of patients with PD compared to controls: None of these genes though appeared as DEGs in any of the comparisons of this study. Pathways that were dysregulated in PD hippocampus included oxidative stress, insulin signaling pathway, neurotransmitter receptors, synaptic and vesicular pathways, protein degradation, and amyloid precursor processing, showing some overlap with the pathways observed in this study. Poor overlap of DEGs, but at least partial overlap of pathways, between animal models and human disease has been highlighted already for inflammatory diseases ^57,58^, AD ^59^, and PD ^37^. It should also be noted that most transcriptional profiling studies of human brain are performed on post-mortem collected tissues, which are, in contrast to the mouse model tissues of this study, mostly reflective of the latest disease stages. The study by Grunblatt et al. also did not specifically highlight sex-specific gene alterations. Thus, further omics profiling studies on PD hippocampus, ideally longitudinal ones, as has been done for PD Substantia Nigra ^60^, are warranted.

Differences in hippocampal transcriptomic profiles between sexes could be explained by intrinsic differences in hippocampal structure between males and females ^61^, and differences in hormonal regulation. Oestrogens and androgens have both been reported to regulate hippocampal synaptic plasticity and neurogenesis ^62,63^, and to modulate learning and memory ^64–66^. Further studies are needed to unravel the role of sex hormones on hippocampal genes changes involved in PD-associated cognitive decline.

Clinically, sex differences have been reported for PD-associated cognitive impairment ^67^. Men and women exhibit different disease courses and outcomes. Men, at least in some ethnicities, are more commonly affected by PD and its progression to PDD, while women are more frequently diagnosed with DLB in certain age ranges. Men often have better performance on certain cognitive tests compared to women. Women with PD tend to be younger at their last clinical visit and may have a shorter disease course, especially in PDD cases. One study confirmed dementia to have an overall higher prevalence in men, although prevalence in women increased after 65 years of age and became equal to that of men from 80 years of age on ^68^.

An informative recent study investigated how PD affects brain structure and function with aging, focusing on differences between male and female patients ^69^. Using neuroimaging techniques, the authors estimated brain age with the “Brain-Predicted Age Difference” (Brain-PAD), which measures the gap between predicted brain age and actual age. A positive Brain-PAD means the brain appears older, while a negative Brain-PAD suggests a younger brain. Men with PD showed higher Brain-PAD, indicating more pronounced brain aging, which correlated with worse cognitive performance. In women, cognitive decline was more related to actual age. These findings further highlight the need for sex-specific approaches in understanding and treating PD-associated cognitive decline.

Taken together, all the studies, including the present one, underscore the importance of considering age and sex in the preclinical and clinical investigation, prognosis, diagnosis, management, and treatment of PD-associated cognitive impairment.

## Materials and Methods

### Animals

#### Transgenic mouse line

The transgenic mouse line BAC-Tg3(SNCA*E46K), which overexpresses the E46K mutated human αSyn was used. This line, also known as Tg(SNCA*E46K)3Elan/J, or“Line 3”, was generated by Elan Pharmaceuticals (USA) ^70^. The longitudinal transcriptional profile of the ventral midbrain and age-dependent striatal neurodegeneration in this model have been reported ^31^. Mice were kept on a C57Bl6/J x DBA/2J hybrid background as described ^31^. They were bred in specific pathogen-free conditions (SPF), with *ad libitum* access to water and food, and under a regular 12h day-night cycle. For each age group (3, 6, 8, and 14 months), 6 to 10 heterozygote *SNCA* transgene carriers (BAC-Tg3(SNCA*E46K)) males, 6 to 10 BAC-Tg3(SNCA*E46K) females, and an equal number of wild-type littermates (WT) were used for histology and protein biochemistry. For bulk RNAseq of hippocampus, 4 animals per group were used. Mice for each experimental group were randomly selected out of at least 3 different litters.

The experiments conformed to the European Union directive 2010/63/EU and the principles of replacement, reduction and refinement (https://www.nc3rs.org.uk/the-3rs). They had received ethical approval by the appropriate Luxembourg governmental entities and were authorized under the project TermProc-UL (LUPA_2020/23).

#### Genotyping

DNA was isolated from ear biopsies using the Extracta^TM^ DNA Prep for PCR-Tissue kit (QuantaBio, #95091) according to manufacturer’s instructions. Briefly, 30 µl of Extraction Reagent were added and the samples were incubated for 30 minutes at 95°C. After incubation, an equal volume of Stabilization Buffer was added and DNA was kept on ice for immediate use or stored at-20°C. For genotyping, the KAPA PROBE FAST UNI genotyping kit (Sigma Aldrich, # KK4702) was used according to manufacturer’s instructions. Each reaction consisted of 2 µl of DNA, 12.5μl of 2xKAPA2G Fast Hot Start Genotyping Mix, 1.25µl of forward and reverse primer at a concentration of 10 µM (5’-3’: forward GAT-TTC-CTT-CTT-TGT-CTC-CTA-TAG-CAC-TGG, reverse GAA-GCA-GGT-ATT-ACT-GGC-AGA-TGA-GGC), and water up to 23 µl. The PCR products were separated by gel electrophoresis and visualised under UV light using a Biometra BioDocAnalyze (Westburg).

#### Euthanasia and tissue collection

Mice were deeply anaesthetised using a mixture of Ketamine (150 mg/kg) and Medetomidine (1 mg/kg) via intraperitoneal injection, and transcardially perfused with PBS to remove the blood. After euthanasia, brains were extracted and split longitudinally in two hemibrains. One hemibrain was fixed in 4% paraformaldehyde for 48h at 4°C, cut into 50 µm thick free-floating sections using a Leica vibratome VT1000 (Wetzlar), and the sections were stored at-20°C in a cryoprotectant medium (1% polyvinyl pyrrolidone, 50% ethylene glycol in PBS) until immunohistochemical staining. The other hemibrain was dissected into regions of interest, which were snap frozen in dry ice and stored at-80°C until use for molecular biology or protein analyses.

### Protein analysis

#### Protein extraction

For protein extraction, the hippocampus was homogenised in 400 µl of lysis buffer (10 mM Tris, 0.32 M sucrose, 2 mM EDTA, 1 mM sodium orthovanadate (Sigma), and antiprotease cocktail (Roche)) using a dounce homogenizer, followed by homogenisation through a 27G needle for 10 times. Protein extracts were snap frozen in dry ice and stored at-80°C until use. Protein extracts’ concentration was determined by Bradford’s method assay (Protein Assay Dye Reagent Concentrate, BioRad, #5000006).

#### Western Blot

Protein analyses were performed as previously described ^37^. Membranes were incubated with primary antibodies (rabbit polyclonal anti-pan-αSyn antibody (S3062 Sigma-Aldrich, AB_477506, 1:1000); mouse monoclonal anti-human-αSyn (clone Syn211, S5566 Sigma-Aldrich, AB_261518, 1:1000); rabbit polyclonal anti-beta-Actin antibody (ab8227 Abcam, AB_2305186, 1:1000)) overnight at 4°C, and with the appropriate secondary antibodies (IRDyeR 800CW donkey anti-rabbit, AB_2715510/ AB_621848, 1:10000 (LI-COR Biosciences); IRDyeR 680LT donkey anti-mouse, (AB_2814906/AB_10715072, 1:10000 (LI-COR Biosciences)) for 1h at RT. There is no available working antibody specifically recognising murine αSyn, so levels thereof were inferred from the level of total αSyn in WT mice, and from the difference between total αSyn and human αSyn in BAC-Tg3(SNCA*E46K) mice. Antibody signal was detected using the Odyssey® CLx Infrared Imaging System (LI-COR Biosciences). Alpha-synuclein protein concentration was quantified in relation to beta-Actin using ImageJ.

#### Immunofluorescence staining on free-floating brain sections

Immunofluorescence staining for human and total αSyn was performed on free-floating brain sections as previously described ^31,71^. A mouse monoclonal anti-human-αSyn (clone Syn211, S5566, Sigma-Aldrich, AB_261518, 1:1000) antibody, a rabbit polyclonal anti-total (pan)-αSyn (S3062 Sigma-Aldrich, AB_477506, 1:1000) and appropriate Alexa Fluor® secondary antibodies were used. The sections were imaged with a Zeiss AxioImager Z1 upright microscope equipped with a“Colibri” LED system and a Zeiss MR R3 digital camera under the control of the Zeiss Blue Vision software. Tiled (3×3 or 3×4) pictures of the whole hippocampus were taken at 10X magnification and then converted into single Tiff files.

#### Synaptophysin quantification

Synaptophysin levels were assessed in the cortical II-V layers and in the hippocampal pyramidal layer, as previously described ^36,72^. Two randomly selected sections per mouse were stained using a mouse monoclonal anti-synaptophysin antibody (NCL-L-SYNAP-299, Leica Biosystems, AB_564017, 1:300)). After immunostaining, slides with brain sections were coded, randomised, and imaged with a Zeiss LSM 710 laser scanning confocal microscope, running a Zeiss Blue Vision software. For each mouse, a total of 6 (3 per section) cortical images and of 4 (two per section) hippocampal images were obtained. Average pixel intensity of synaptophysin staining was calculated using NIH image, and the values were averaged for each region/mouse. The codes of the slides were broken only after analysis was complete.

### RNA analyses

#### RNA extraction

RNA was extracted from frozen hippocampi with the RNeasy Plus Universal Mini kit (Qiagen, #73404) according to manufacturer’s instructions and as previously described ^37^. RNA was kept on ice for immediate use or stored at-80°C. RNA quality and concentration were measured using a NanoDrop^TM^ 2000 (ThermoFischer Scientific) and an Agilent 2100 Bioanalyzer.

#### RT-qPCR

For retro-transcription, the SuperScript^TM^ III RT reverse transcriptase (Invitrogen, #18080093) was used according to the manufacturer’s instructions and as previously described ^31,37^.

The Real Time (RT)-qPCR reaction was performed as previously described ^31^ using the following primers: murine *Snca* (forward CGC-ACC-TCC-AAC-CAA-CCC-G, reverse TGA-TTT-GTC-AGC-GCC-TCT-CCC); human *SNCA* (forward AAG-AGG-GTG-TTC-TCT-ATG-TAG-GC, reverse GCT-CCT-CCA-ACA-TTT-GTC-ACT-T); *GAPDH* (forward TGC-GAC-TTC-AAC-AGC-AACTC, reverse CTT-GCT-CAG-TGT-CCT-TGC-TG). Expression of each target gene was normalised to *GAPDH* (ΔCt) and expressed as fold change (2^(-ΔCt)).

#### mRNA sequencing

RNA sequencing was performed by the sequencing platform of the Luxembourg Centre for Systems Biomedicine (LCSB) at the University of Luxembourg, using a total of 1 µg of RNA. Library preparation was performed using the Illumina Stranded mRNA Prep kit, and library quantification was performed using a QubitTM 4 Fluorometer (Invitrogen). The NextSeq2000 (Illumina) was used for 2 x 50 bp paired end sequencing with a sequencing depth of ∼24 million reads per sample.

Raw sequencing data obtained as fastq files were quality-checked and aligned as previously described ^30^ using STAR v2.7.9a ^73^ against a custom-built genome based on the GRCm39 mouse assembly and the human *SNCA* gene locus. Raw expression values were determined based on Ensembl gene annotations v104 and imported into *DESeq2* ^74^ under *R*. Genes with less than 20 mean reads across samples were excluded from subsequent analyses, leaving ∼18.500. Unwanted variation between samples was minimised with surrogate variable analysis (*sva*, v3.44). Differential genes were identified using a cut-off of Benjamini-Hochberg adjusted *p*-value ≤ 0.05. Relative gene expression was calculated as normalised Reads Per Kilobase per Million total reads (nRPKMs) ^75^. Cell type composition across samples was estimated based on cell type-specific reference data^41^.

#### Pathway analysis

Pathway analysis was performed using QIAGEN IPA (Release Dec 15, 2023, https://digitalinsights.qiagen.com/IPA) using default settings, and with the analysis limited to the mouse genome.

To get an overarching view of the aging and disease processes in the hippocampus of male and female BAC-Tg3(SNCA*E46K) mice and their wildtype littermates, the individual pathways obtained through IPA were manually grouped into high-order functional biological clusters. Publicly available online tools (Google, Wikipedia, ChatGPT 3.5) were used to identify abbreviations used by the IPA software, and to attribute the pathways to functional clusters when the name of a pathway or its relevance for the CNS weren’t obvious (Suppl. Table 1 and 2). A number of pathways, being quite generic, were not attributable to high level functions. Then, for each pairwise comparison of hippocampal gene expression profiles (sex-, age-or genotype-dependent), a“●” was added to a cluster table cell each time a pathway attributed to that cluster appeared in the IPA analysis of that comparison (Table 1).

## Declaration of generative AI and AI-assisted technologies in the writing process

During the preparation of this work the authors used ChatGPT 3.5 in order to edit parts of the Results’ section, and help with contextualising the biological significance of IPA pathways. After using this tool/service, the authors reviewed and edited the content as needed, and take full responsibility for the content of the published article.

## Data availability

All RNA-seq data have been uploaded to GEO and are available under the accession number GSE271598. All other original data are available upon request.

## Abbreviations

6-OHDA: 6-hydroxydopamine
αSyn: alpha-synuclein
APBA3: Amyloid Beta Precursor Protein-Binding, Family A, Member 3
APP: Amyloid Beta Precursor Protein
BAC: Bacterial artificial chromosome
BBB: Blood brain barrier
Brain-PAD: Brain-predicted age difference
CA1: Cornu Ammonis 1
CA3: Cornu Ammonis 3
CAS: Common age signature
CHRNA6: Cholinergic Receptor, Nicotinic, Alpha Polypeptide 6
CNS: Central nervous system
CREB: cAMP Response Element-Binding Protein
Ddx3x: DEAD-Box Helicase 3 X-Linked
DEGs: Differentially expressed genes
DLB: Dementia with Lewy bodies
DNA: Deoxyribonucleic acid
ECM: Extracellular matrix
GABA: Gamma-aminobutyric acid
GO: Gene ontology
GP6: Glycoprotein VI Platelet
GSTM1: Glutathione S-Transferase M1
GTPases: guanosine triphosphate binding proteins
HET: Heterozygous
IPA: Ingenuity Pathway Analysis
Kdm5c: Lysine Demethylase 5C
Kdm5d: Lysine Demethylase 5D
log_10_p: logarithm base 10 of the p-value
MAPK: Mitogen-activated protein kinase
MCI: mild cognitive impairment
MPTP: 1-methyl-4-phenyl-1,2,3,6-tetrahydropyridine
NMDA: N-methyl-D-aspartate
PCA: Principal component analysis
PD: Parkinson’s Disease
PDD: Parkinson’s Disease Dementia
RNA: Ribonucleic acid
RNAseq: RNA sequencing
RT-qPCR: Quantitative reverse transcription polymerase chain reaction
SNCA: alpha-synuclein, human
Snca: alpha-synuclein, mouse
Tg: transgenic
VPS35: Vacuolar Protein Sorting 35
WT: wild-type littermate
Xist: X-inactive specific transcript

## Author contributions: *Alessia Sciortino*

Investigation, Formal analysis, Visualisation, Writing - Original Draft. ***Thomas Hentrich:*** Formal analysis, Data curation, Visualisation, Writing - Review & editing. ***Sergio Helgueta:*** Formal Analysis, Data curation. ***Pierre Garcia:*** Investigation. ***Kristopher J Schmit:*** Investigation. ***Mélanie Thomas:*** Investigation. ***Rashi Halder:*** Investigation. ***Djalil Coowar***: Conceptualisation, Supervision. ***Michel Mittelbronn:*** Funding acquisition, Supervision. ***Lasse Sinkkonen:*** Conceptualisation. ***Julia Schultze-Hentrich:*** Conceptualisation. ***Manuel Buttini:*** Conceptualisation, Supervision, Funding acquisition, Investigation, Formal Analysis, Writing-Review & editing.

## Acknowledgments

The authors thank the animal facility support team, and the pre-publication-check team at the University of Luxembourg, for help and support. They also thank Janine Schulze for help with RNAseq.

