## Supplemental Tables for "Transcriptional Dysregulation in the Hippocampus of a murine model for Parkinson’s Disease Cognition Impairment is Driven by Sex, Age, and Alpha-synuclein overexpression"

**Suppl. Table 1**: IPA abbreviations. Public online resources (Google, Pubmed, Wikipedia) were used to find the whole names of abbreviations used by IPA.

| **Abbreviation** | **Full name** | **Most relevant function(s), including in the CNS if known** |
|---|---|---|
| ATF6 | Activating transcription factor 6 | Induces the endoplasmatic reticulum unfolded protein stress response |
| BBsome | Bardel-Bield Syndrome 8 protein complex | Orchestrates cilia function |
| CREB | cAMP response element-binding protein | Transcription factor |
| CXCR4 | CXC motive chemokine receptor 4 | Receptor for CXCL12 |
| DAG/IP3 | Diacylglycerol/inositol 3-phosphate | Intracellular calcium release; neuronal survival/death and synaptic plasticity |
| EIF2AK4 (GCN2) | Eukaryotic translation initiation factor 2 alpha kinase 4 (General Control non-derepressible 2) | Downregulates protein synthesis in response to cellular stress |
| ERBB | family of receptor tyrosine kinases | Involved in multitude of neuronal functions |
| ERBB 4 | Erb-B2 Receptor Tyrosine Kinase 4 | Tyrosine kinase receptor, multitude of neuronal functions |
| GAL | Galanin | Modulatory neuropeptide |
| GP1B-IX-V | Glycoprotein 1b-IX-V complex membrane receptor | Role in platelet adhesion/thrpmsbosi; orle in rain unclear |
| GP6 | Glycoprotein 6 | Receptor for collagen on platelets, induces platelet aggregation, function in CNS unclear |
| GRNH | Gonadotropin-releasing hormone | Synaptic plasticity and neuroprotection in the hippocampus |
| HEY1 | Hairy enhancer of Split-related with YRPW motif protein 1 | Mediator of NOTCH1, transcriptional repressor |
| HIFa | Hypoxia-inducible factor 1 | Response to hypoxia/cellular stress |
| HMOX1 | Heme oxygenase 1 | Heme degradation, anti-oxidant and anti-inflammatory |
| HOTAIR | HOX (homeobox) transcript antisense RNA | Role in gene expression regulation through histone modification; induced by cellular stress |
| IGF and IGFBPs | Insulin-like growth factor and IGF-binding proteins | |
| IL10 | Interleukin 10 | Inflammatory cytokine |
| Il17A | Interleukin 17A | Anti-inflammatory cytokine |

| | | |
|---|---|---|
| LXR-RXR | Liver X receptor- retinoic R receptor | Form heterodimers to regulate gene expression, regulating cholesterol metabolism and acting anti-inflammatory |
| MECP2 | Methyl CpG binding protein | Maintenance of mature neurons |
| MET | Mesenchymal epithelial transition factor | Neuroprotection/-regeneration, synaptic plasticity |
| NFAT | Nuclear Factor of Activated T cells | Modulation of immune response, differentiation of T cells |
| NMD | Nonsense-Mediated mRNA Decay | Quality control mechanism in eukaryotic cells to eliminate aberrant, incomplete mRNA transcripts; enhanced during cellular stress |
| NOD 1/2 | Nucleotide-binding oligomerization domain containing protein 1/2 | Immune-related pattern recognition receptor; induce pro-inflammatory response |
| PDGF | Platelet-derived growth factor | Various functions and sources in the brain |
| PIP2 | Phosphatidylinositol 4,5-biphosphate | Component of cell membranes and signaling molecule |
| PPAR | Peroxisome-proliferator active receptor | Glucose and fatty acid metabolism regulation; role in neuroinflammation |
| RANK | Receptor activation of nuclear factor kappa B | Roles in bone remodeling and immune function |
| RHO GTPases | Ras homolog gene family member guanosine triphosphatase hydrolase | Transduce signals from extracellular stimuli to intracellular signaling pathways for multiple functions |
| ROBO SLIT | Roundabout (receptor) and its Slit guidance ligand | Axonal guidance in the CNS |
| S100 | "Soluble in 100% ammonium sulfate solution" family of proteins | Calcium-binding proteins with multiple functions |
| TNFR2 | Tumor necrosis factor receptor 2 | Multiple functions in the brain |
| TP53 | Tumor protein 53 | Intranuclear protein controlling cell division and death, and transcription of metabolic genes |
| TR/RXR | Thyroid hormone receptor and retinoic X receptor | Regulation of neuronal differentiation, neurogenesis, synaptic plasticity |
| TSP1 | Thrombospondin 1 | Cell/Cell and Cell-Extracellular Matrix interactions, collagen homeostasis |
| TRAP | Tumor necrosis factor receptor associated protein | |
| UPR | Unfolded protein response | Stress response |
| VEGF | Vascular endothelial growth factor | Stimulation of blood vessel formation |

**Suppl. Table 2**: Grouping of IPA pathways into functional clusters.

| Functional clusters | IPA pathway | Sex-dependent | Age-dependent | | Genotype-dependent | Remarks/justification |
|---|---|---|---|---|---|---|
| | | | males | females | | |
| **Neuronal function**<br>***Synapse*** | Synaptogenesis signaling pathway | WT, 6m | TG, 14m-8m | WT, 8m-6m<br>TG, 14m-8m | | |
| | SNARE (Soluble N-ethylmaleimide-sensitive factor activating protein receptor) signaling pathway | WT, 6m | | | | Membrane fusion and neurotransmitter release |
| | GNRH (Gonadotropin-releasing hormone) signaling | TG, 3m | TG, 14m-8m | | | Regulates synapse spine density via estrogen synthesis |
| ***Axon*** | Axonal guidance signaling | | TG, 14m-8m | WT, 8m-6m<br>TG, 14m-8m | M, 3m<br>F, 6m | |
| | ROBO SLIT (Roundabout receptor and its Slit guidance ligand) signaling pathway | WT, 14m | | | | Controls axonal guidance in CNS |
| | Semaphorin neuronal repulsive signaling pathway | | | | F, 6m | Role in axonal and growth cone guidance |
| ***Neuronal activity /plasticity*** | Sleep REM signaling pathway | WT, 14m<br>TG, 6m | | | | Involves neurotransmitters, neuropeptides, neurohormones |
| | Sleep NREM signaling pathway | | | | F, 6m | |
| | Signaling by ERBB4 (Erb-B2 Receptor Tyrosine Kinase 4) | | WT, 8m-6m | | | Neuronal development, synaptic plasticity, myelination |
| | Signaling by MET (Mesenchymal epithelial transition factor) | | | WT, 6m-3m | | Regulation of neural circuits, synapses, neuron-glia interaction |
| | Neuropathic pain signaling on dorsal horns | | | WT, 8m-6m | | |
| | Growth hormone signaling | | | WT, 14m-8m<br>TG, 8m-6m | F, 3m | Neurogenesis, synaptic plasticity |
| | Class A/1 (Rhodopsin-like receptors) | | | TG, 6m-3m | M, 14m | In hippocampus, receptors involved mainly in regulation of neuronal and synaptic activity |
| | Breast cancer regulation by Stathmin1 | | | TG, 6m-3m | M, 14m | In hippocampus, stathmin 1 is produced by neurons and regulates their morphology and synaptic plasticity |

| | | | | | | |
|---|---|---|---|---|---|---|
| ***Neuronal gene transcription & regulation*** | NFG (nerve growth factor) stimulated transcription | TG, 3m<br>TG, 14m | | TG,<br>14m-8m | F, 8m | |
| | Corticotropin Releasing Hormone Signaling | TG, 3m | WT,<br>14m-8m | | | In the hippocampus, regulates synaptic plasticity and learning |
| | Transcriptional regulation by MECP2 (Methyl CpG binding protein) | TG, 6m | WT,<br>8m-6m | | | Maintenance of mature neurons |
| | CREB (cAMP response element-binding protein) signaling in neurons | | | | F, 6m | Transcription factor |
| ***Glutamate neurotransmission*** | Glutaminergic receptor signaling pathway (enhanced) | WT, 8m | TG,<br>6m-3m<br>TG,<br>8m-6m | | F, 6m | |
| | Assembly and cell surface presentation of NMDA (N-methyl-D-aspartate) receptors | WT, 8m | | | | |
| | Glutamate receptor signaling | | | | F, 14m | |
| | ERBB (family of receptor tyrosine kinases) signaling | | WT,<br>6m-3m | | | Key factor for glutaminergic circuit assembly and synapse development |
| | Activation of NMDA (N-methyl-D-aspartate) receptors and postsynaptic events | | WT,<br>8m-6m | WT,<br>8m-6m | | |
| ***Other neuropeptides / -transmitters*** | Opioid signaling pathway | TG, 3m | TG,<br>14m-8m | WT,<br>8m-6m<br>TG,<br>14m-8m | F, 6m<br>F ,14m | |
| | GABA (Gamma-aminobutyric acid) receptor activation, GABA receptor signaling | TG, 6m | WT,<br>8m-6m | | F,<br>14m | |
| | Dopamine-DARPP32 feedback in cAMP signaling | | WT,<br>14m-8m | | | |
| | Endocannabinoid neuronal synapse pathway | | | WT,<br>8m-6m | F, 14m | |
| | Serotonin receptor signaling | | | | M, 14m | |
| **Neuroprotection** | Insulin secretion signaling pathway | TG, 3m | | WT,<br>14m-8m | | In the CNS, involved in neuroprotection, synaptic plasticity, memory |
| | TR/RXR (Thyroid hormone receptor and retinoic X receptor) activation | TG, 14m | | | | Regulation of neuronal differentiation, neurogenesis, synaptic plasticity |
| | LXR/RXR (Liver X receptor-retinoic R receptor) activation | | | TG,<br>6m-3m | | Neuroprotection, anti-inflammatory action, |

| | | | | | | |
|---|---|---|---|---|---|---|
| | | | | | | cholesterol metabolism regulation |
| | Stearate biosynthesis | | TG, 8m-6m | | | Neuroprotectant |
| | Seleno-amino-acid metabolism | | TG, 14m-8m | | | Linked to neuroprotection |
| | DHA (Docosahexaenoic acid) signaling | | | | M, 14m | Part of neuronal membrane, role in neuroprotection |
| **Immune function** | Role of NFAT (Nuclear Factor of Activated T cells) in cardiac hypertrophy | WT, 6m | | | | Modulation of immune response (ref), differentiation of T cells |
| | Granzyme A signaling | WT, 8m | | | | |
| | CXCR4 (CXC motive chemokine receptor 4) | TG, 3m | | | | Inflammation and development |
| | Calcium-induced T lymphocyte apoptosis | TG, 3m | | | | |
| | Neutrophil extracellular TRAP (Tumor necrosis factor receptor associated protein) signaling pathway | TG, 3m | TG, 6m-3m | | | |
| | Binding and uptake of ligands by scavenger receptors | TG, 8m | | | | |
| | Immunogenic cell death signaling pathway | TG, 8m | | | | |
| | Leukocyte Extravasation Signaling | | WT, 6m-3m TG, 8m-6m | | | |
| | LPS-stimulated MAPK signaling | | WT, 6m-3m | | | |
| | RANK (Receptor activation of nuclear factor kappa B) signaling | | WT, 6m-3m | | | Roles in bone remodeling and immune function |
| | Fcg- receptor-mediated phagocytosis in macrophages | | WT, 6m-3m | | | |
| | Neuroinflammation signaling pathway | | TG, 8m-6m | | | |
| | Arachidonic acid metabolism | | TG, 8m-6m | | | |
| | Binding and uptake of ligands by scavenger receptors | | | WT, 6m-3m | | |
| | NOD1/2 (Nucleotide-binding oligomerization domain containing protein 1/2) signaling pathway | | | WT, 14m-8m | | Immune-related pattern recognition receptor; induce pro-inflammatory response |
| | IL-12 (Interleukin 12) signaling and production in macrophages | | | WT, 14m-8m | | |
| | Granulocyte adhesion and diapedesis | | | TG, 6m-3m | | |

| | | | | | | |
|---|---|---|---|---|---|---|
| | Pathogen-induced cytokine storm signaling pathway | | | TG, 8m-6m | | |
| | Neutrophil degranulation | | | | M, 3m | |
| | Role of JAK-family kinases in IL-6-type cytokine signaling | | | | M, 6m | |
| | Phagosome formation | | | | M, 14m | |
| | Neutrophil extracellular TRAP (Tumor necrosis factor receptor associated protein) signaling pathway | | | | M, 14m<br>F, 3m | |
| | Macrophage alternative activation signaling pathway | | | | F, 8m | |
| | IL-17A (Interleukin 17A) signaling in fibroblasts | | | | F, 8m | Anti-inflammatory cytokine |
| | IL-10 (Interleukin 10) signaling | | | | F, 8m | Inflammatory cytokine |
| **Astrocyte reactivity** | Hepatic fibrosis/ hepatic stellate cell activation | WT, 3m<br>WT, 14m<br>TG, 6m | WT, 14m-8m | WT, 6m-3m<br>TG, 8m-6m<br>TG, 14m-8m | M, 3m<br>M, 6m<br>F, 3m | Parallels between astrocytes and hepatic stellate cells (e.g. collagen production) |
| | Wound healing signaling pathway | WT, 3m | | | F, 8m | Astrocytes orchestrate wound healing and scar formation in the brain |
| | Sertoli Cell-Sertoli Cell Junction Signaling | | WT, 6m-3m | | | Astrocytes and Sertoli cells have common functionalities (nutritional/ modulatory support, barrier formation) |
| **Stress/antioxidant/ protein degradation & misfolding responses** | Cytoprotection by HMOX1 (Heme oxygenase 1) | WT, 8m | | | | Stress-response gene involved in heme degradation, anti-oxidant and anti-inflammatory responses |
| | Response of EIF2AK4 (GCN2) (Eukaryotic translation initiation factor 2 alpha kinase 4 (General Control non-derepressible 2)) to amino acid deficiency | WT, 8m<br>TG, 14m | TG, 14m-8m | | | EIF2AK4 downregulates protein synthesis in response to cellular stress |
| | HIF alpha (Hypoxia-inducible factor 1) signaling | WT, 14m | | | | Response to hypoxia/cellular stress |
| | HOTAIR (HOX (homeobox) transcript antisense RNA) regulatory pathway | WT, 14m | | | M, 8m | Role in gene expression regulation through histone modification; induced by cellular stress |

| | | | | | | |
|---|---|---|---|---|---|---|
| | ATF6-alpha (Activating transcription factor 6) activates chaperone genes | TG, 8m<br>TG, 14m | | | | Induces the endoplasmatic reticulum unfolded protein stress response |
| | Microautophagy signaling pathway | WT, 3m | | | | |
| | Protein ubiquitination pathway | | | | M, 8m | |
| | Endoplasmatic reticulum stress pathway | TG, 8m | | | | |
| | Unfolded protein response (UPR) | TG, 8m<br>TG, 14m | | WT,<br>14m-8m | | Stress response |
| | UVB-induced MAPK signaling | | WT,<br>6m-3m | | | regulates processes of cell survival, DNA repair, inflammation, and apoptosis in response to stress |
| | Ferroptosis signaling pathway | | WT,<br>8m-6m | | | Form of regulated cell death in response to stress |
| | Nonsense-Mediated Decay (NMD) | | TG,<br>14m-8m | | | Degradation of non-sensical and aberrant RNAs |
| | Cellular response to heat stress | | | WT,<br>14m-8m | | |
| | Acute phase response signaling | | | | F, 8m | |
| **Vascular processes** | Neurovascular coupling signaling pathway | WT, 3m | WT,<br>8m-6m<br>TG,<br>6m-3m | | M, 6m<br>F, 14m | Included by cellular stress |
| | Signaling by VEGF (Vascular endothelial growth factor) | | WT,<br>6m-3m | | | Stimulation of blood vessel formation |
| | VEGF (Vascular endothelial growth factor) family ligand receptor interactions | | | | M, 6m | |
| | NO (nitric oxide) signaling in the cardiovascular system | | WT,<br>14m-8m | | | |
| | Cellular effects of sildenafil | | WT,<br>14m-8m | | | |
| | eNOS (endothelial nitric oxide synthase) signaling | | WT,<br>14m-8m | | | |
| | Atherosclerosis signaling | | TG,<br>8m-6m | | M, 6m | |
| | Apelin liver signaling pathway | | | WT,<br>6m-3m | | Apolin is vasoconstrictive and lowers blood pressure |
| **Collagen related** | Collagen chain trimerization | WT, 3m<br>TG, 3m | TG,<br>6m-3m | WT,<br>6m-3m<br>TG,<br>14m -8m | M, 6m | |
| | Assembly of collagen fibrils | WT, 3m | TG, | WT, | | |

| | | | | | | |
|---|---|---|---|---|---|---|
| | | | 6m-3m | 6m-3m<br>TG,<br>14m-8m | | |
| | Collagen degradation | WT, 3m | TG,<br>6m-3m | WT,<br>6m-3m | M, 6m | |
| | Collagen biosynthesis and modifying enzymes | WT, 3m<br>WT, 14m | TG,<br>6m-3m | WT,<br>6m-3m | M, 6m | |
| | GP6 (Glycoprotein 6) receptor signaling, GP6 signaling pathway | WT, 3m | TG,<br>6m-3m | | F, 3m | Receptor for collagen on platelets, induces platelet aggregation, function in CNS unclear |
| | Inhibition of angiogenesis by TSP1 (Thrombospondin 1) | WT, 14m | | | | Cell/Cell and Cell-Extracellular Matrix interactions, collagen homeostasis |
| **Extracellular matrix (ECM)** | ECM organization | WT, 3m | | TG,<br>8m-6m | F, 14m | |
| | Elastic fiber formation | | WT,<br>8m-6m | | | Elastin is part of the ECM |
| | Degradation of the ECM | | WT,<br>8m-6m | | | |
| | Integrin cell surface interactions | | TG,<br>6m-3m | TG,<br>8m-6m<br>TG,<br>14m-8m | M, 6m | Integrins are cell surface receptors interacting with ECM |
| **Calcium signaling And regulation** | Calcium signaling | WT, 6m<br>TG, 6m | WT,<br>8m-6m | WT,<br>8m-6m<br>TG,<br>8m-6m | F, 6m | |
| | Calcium transport 1 | TG, 6m | | | | |
| | Response to elevated platelet cytosolic Ca2+ | | | | F, 3m | |
| | S100 ("Soluble in 100% ammonium sulfate solution" family of proteins) family signaling pathway | | TG,<br>8m-6m | | M, 14m | Calcium-binding proteins with multiple functions |
| | DAG and IP3 (Diacylglycerol/inositol 3-phosphate) signaling | | | WT,<br>8m-6m | | Intracellular calcium release; neuronal survival/death and synaptic plasticity |
| **Metabolic** | Tumor protein 53 (TP53) regulates metabolic genes | WT, 8m | | | | Intranuclear protein controlling cell division &death, and transcription of metabolic genes |
| | Glycogen metabolism | TG, 14m | | | | |
| | PPAR (Peroxisome-proliferator active receptor) signaling | | | WT,<br>14m-8m | | Glucose and fatty acid metabolism regulation; role in neuroinflammation |
| **Myelin** | Myelination signaling pathway | WT, 6m | | | | |

| | | | | | | |
|---|---|---|---|---|---|---|
| **Mitochondria** | Oxidative phosphorylation | WT, 8m | | | | |
| | Electron transport ATPsyn | WT, 8m | | | | |
| | Mitochondrial dysfunction | WT, 8m | | | | |
| | tRNA processing in the mitochondrion | | WT, 14m-8m | | | |
| | Fatty acid b-oxidation | | TG, 8m-6m | | | |
| **Generic/ unclassifiable** | Role of osteoclasts in rheumatoid arthritis signaling pathway | WT, 3m | | | M, 8m | |
| | Role of chondrocytes in rheumatoid arthritis signaling pathway | | | | F, 8m | |
| | Dilated cardiomyopathy signaling pathway | WT, 6m | | WT, 8m-6m TG, 8m-6m | | In the CNS, neurons are the most similar to cardiocytes because of the electrical stimulability |
| | Cardiac hypertrophy signaling | | | | F, 6m | |
| | Cardiac conduction | WT, 6m | | | | |
| | G-protein coupled receptor signaling | WT, 6m TG, 6m | | | M, 3m F, 6m | |
| | Potassium channels | WT, 6m | | WT, 14m-8m | F, 14m | |
| | Iron homeostasis signaling pathway | WT, 14m | | | | |
| | Glioblastoma multiform signaling | WT, 14m | | | | |
| | Pulmonary fibrosis idiopathic signaling pathway | TG, 3m | | TG, 14m-8m | | TGFbeta1/Smad 3 and Wnt/beta-catenin signaling |
| | Pulmonary healing signaling pathway | | | | M, 6m | |
| | Cardiac conduction | TG, 6m | | | | |
| | Ion channel transport | TG, 6m | | | | |
| | Factors promoting cardiogenesis in vertebrates | TG, 6m | | | | Cells most similar to cardiocytes in the CNS are neurons, which are also electrically stimulatable |
| | Aldosterone signaling in epithelial cells | TG, 8m | | | | Vascular? BBB? |
| | GP1B-IX-V (Glycoprotein 1b-IX-V complex membrane receptor) activation signaling | TG, 8m | | | | Role in platelet adhesion and aggregation, role in CNS unclear |
| | Platelet aggregation (plug formation) | TG, 8m | | | | |
| | Platelet adhesion to exposed collagen | TG, 8m | | | | |
| | Intrinsic prothrombin activation pathway | | | WT, 6m-3m TG, 6m-3m | | |

| | | | | | | |
|---|---|---|---|---|---|---|
| | Platelet homeostasis | | | | M, 8m<br>F, 14m | |
| | O-linked glycosylation | TG, 14m | | | | |
| | Major pathway of rRNA processing in the nucleus and cytosol | TG, 14m | | | | |
| | Anti proliferative role of somatostatin receptor 2 | TG, 14m | | | | |
| | Eukaryotic translation initiation | TG, 14m | | | | |
| | Signaling by Rho Family GTPases (Ras homolog gene family member guanosine triphosphatase hydrolases) | | WT, 6m-3m | TG, 14m-8m | | Transduce signals from extracellular stimuli to intracellular signaling pathways for multiple functions |
| | Renin-Angiotensin Signaling | | WT, 6m-3m | | | |
| | Cardiomyocyte differentiation via BMP receptors | | WT, 8m-6m | | | |
| | HEY1 (Hairy enhancer of Split-related with YRPW motif protein 1) signaling pathway | | WT, 8m-6m | | | Mediator of NOTCH1, transcriptional repressor |
| | rRNA processing | | WT, 14m-8m | TG, 6m-3m | | |
| | Circadian rhythm signaling | | WT, 14m-8m<br>TG, 14m-8m | WT, 8m-6m<br>WT, 14m-8m | M, 8m | |
| | White adipose tissue browning pathway | | WT, 14m-8m | | F, 6m | |
| | Effect of PiP2 (Phosphatidylinositol 4,5-biphosphate) hydrolysis | | TG, 6m-3m | | | Component of cell membranes and signaling molecule |
| | Virus entry via endocytic pathways | | TG, 8m-6m | | | |
| | Eukaryotic translation elongation | | TG, 14m-8m | | | |
| | Eukaryotic translation termination | | TG, 14m-8m | | | |
| | Syndecan interactions | | | WT, 6m-3m | | |
| | Erythrocytes take up oxygen and release carbon dioxide | | | WT, 14m-8m | | |
| | Signaling by PDGF (Platelet-derived growth factor) | | | TG, 6m-3m | F, 14m | Various functions and sources in the brain |
| | Post-translational protein phosphorylation | | | TG, 6m-3m | | |
| | Keratinization | | | TG, 6m-3m | | Keratin protein is expressed in some brain compartments (eg choroid plexus). |
| | Regulation of IGF (Insulin-like growth factor) | | | TG, 6m-3m | F, 8m | |

| | | | | | | |
|---|---|---|---|---|---|---|
| | transport and uptake by IGFBPs (IGF-binding proteins) | | | | | |
| | Protein kinase A signaling | | | TG, 8m-6m | | |
| | Tumor microenvironment pathway | | | TG, 8m-6m | M, 8m | |
| | Transport of bile salts and organic acids, metal ions & amine comp. | | | | M, 8m | |
| | RGO GTPases activate formins | | | | M, 8m | Actin reorganization, cell motility |
| | Mitotic prometaphase | | | | M, 8m | |
| | Sperm motility | | | | M, 8m | Cell motility |
| | G alpha (q) signaling events | | | | M, 14m | |
| | Ga1 signaling | | | | F, 14m | |
| | GAP junction signaling | | | | M, 14m | |
| | BBSome (Bardel-Bield Syndrome 8 protein complex) signaling pathway | | | | M, 14m | Orchestrates cilia function |
| | Post-translational protein phosphorylation | | | | F, 8m | |
| | TNFR2 (Tumor necrosis factor receptor 2) signaling | | | | F, 8m | Multiple functions in the brain |

Public online resources (Google, Wikipedia, Pubmed, ChatGPT 3.5) were used to decipher biological functions of IPA pathways when it was not immediately clear from their naming by the IPA software, and were assigned to functional clusters. Table shows functional clusters (column 1) into which IPA pathways (column 2) were manually grouped. The cluster “Neuronal function” was further subdivided into functional neuronal compartments or activities.  Next 3 columns show the comparisons of gene expression profiles (sex-dependent: column 3, age-dependent: column 4, further divided into males and females, and genotype-dependent: column 5) in which each of listed IPA pathway emerged as significantly changed. (WT = wildtype mice, TG = heterozygous BAC-Tg3(SNCA*E46K) mice; M = males, F = females). Last column shows biological functions of IPA pathways as collected with public online tools.
