## Supplemental Figures for "Transcriptional Dysregulation in the Hippocampus of a murine model for Parkinson’s Disease Cognition Impairment is Driven by Sex, Age, and Alpha-synuclein overexpression"

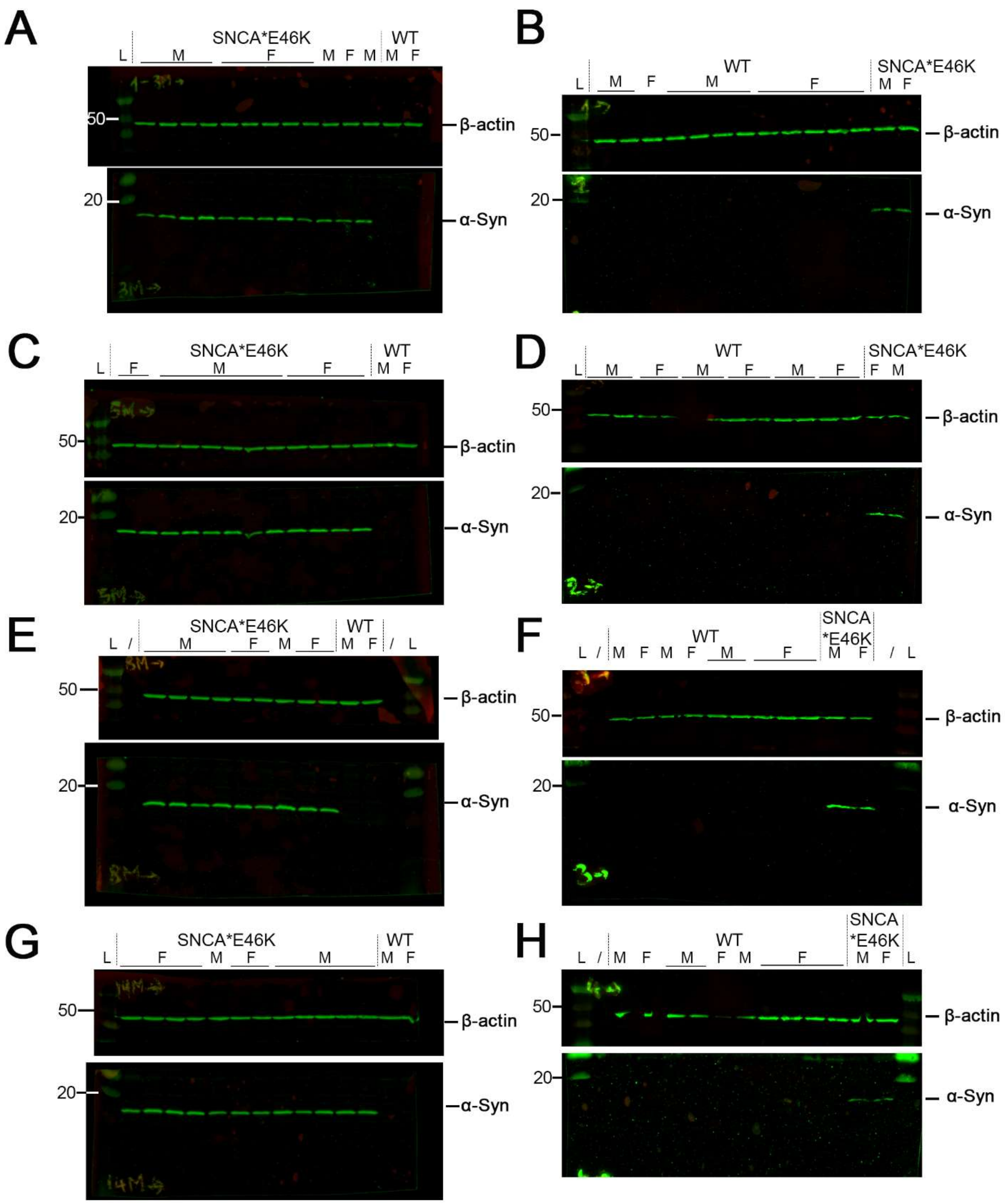


**Suppl. Fig. 1 Western Blot raw images of human alpha-synuclein protein** (h-αSyn, 14kDa) for **A-B)** 3-month- old mice, **C-D)** 5-month-old mice, **E-F)** 8-month-old mice, and **F-G)** 14-month-old mice. β-actin (42kDa) was used as a loading control. Nitrocellulose membranes were cut after transfer, incubated with appropriate primary and secondary antibodies, and imaged in parallel. L = Precision Plus Protein Dual Color Standards. Only the green channel was used, thus causing inefficient visualisation of protein standards in some images.

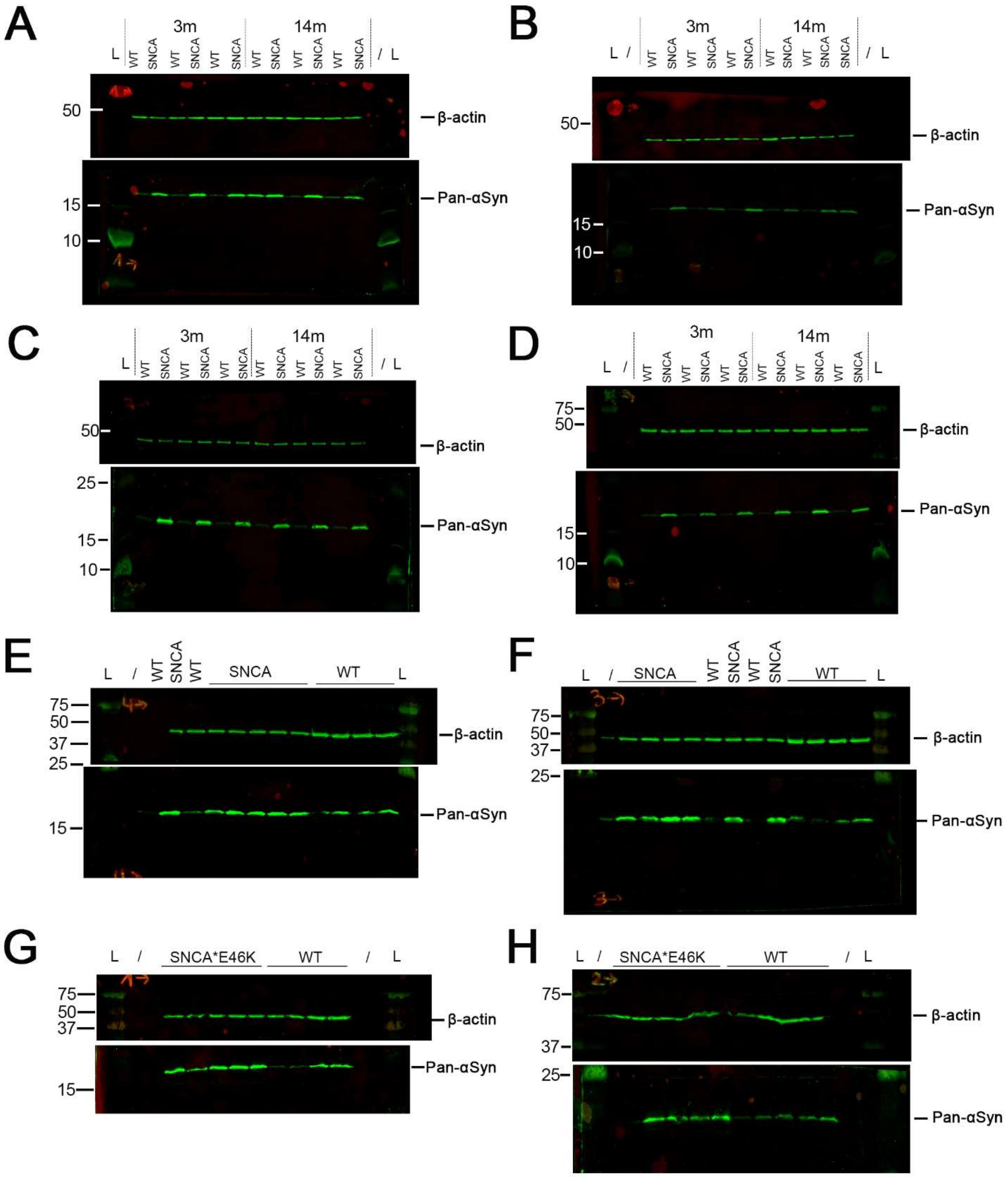


**Suppl. Fig. 2 Western Blot raw images of total alpha-synuclein protein** (Pan-αSyn, 14kDa) for **A-B)** 3- and 14-month-old male mice, **C-D)** 3- and 14-month-old female mice, **E)** 5-month-old male mice, **F)** 5-month-old female mice, **G)** 8-month-old male mice, and **H)** 8-month-old female mice. β-actin (42kDa) was used as a loading control. Nitrocellulose membranes were cut after transfer, incubated with appropriate primary and secondary antibodies, and imaged in parallel. L = Precision Plus Protein Dual Color Standards. Only the green channel was used, thus causing inefficient visualisation of protein standards in some images.

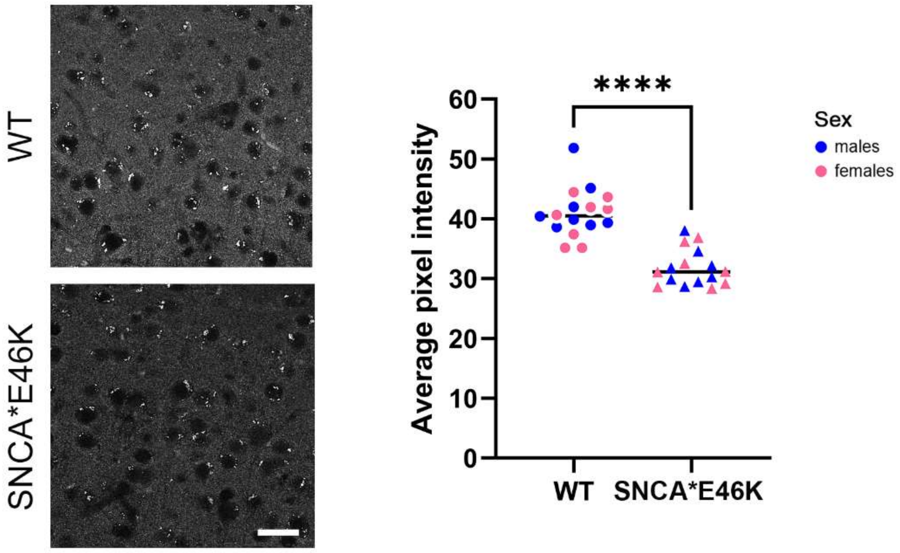


**Suppl. Fig. 3 SNCA overexpression leads to loss of synaptophysin in the cortex of BAC-Tg3(SNCA*E46K) mice.** Representative images for Synaptophysin (SYP) immunostaining (Scale bar: 50 µm) and quantification of SYP staining intensity in the cortex of BAC-Tg3(SNCA*E46K) and WT mice. No difference between male and female mice of the same genotypes was detected. N=6-8/group, **** = $p < 0.0001$ (Student’s t).

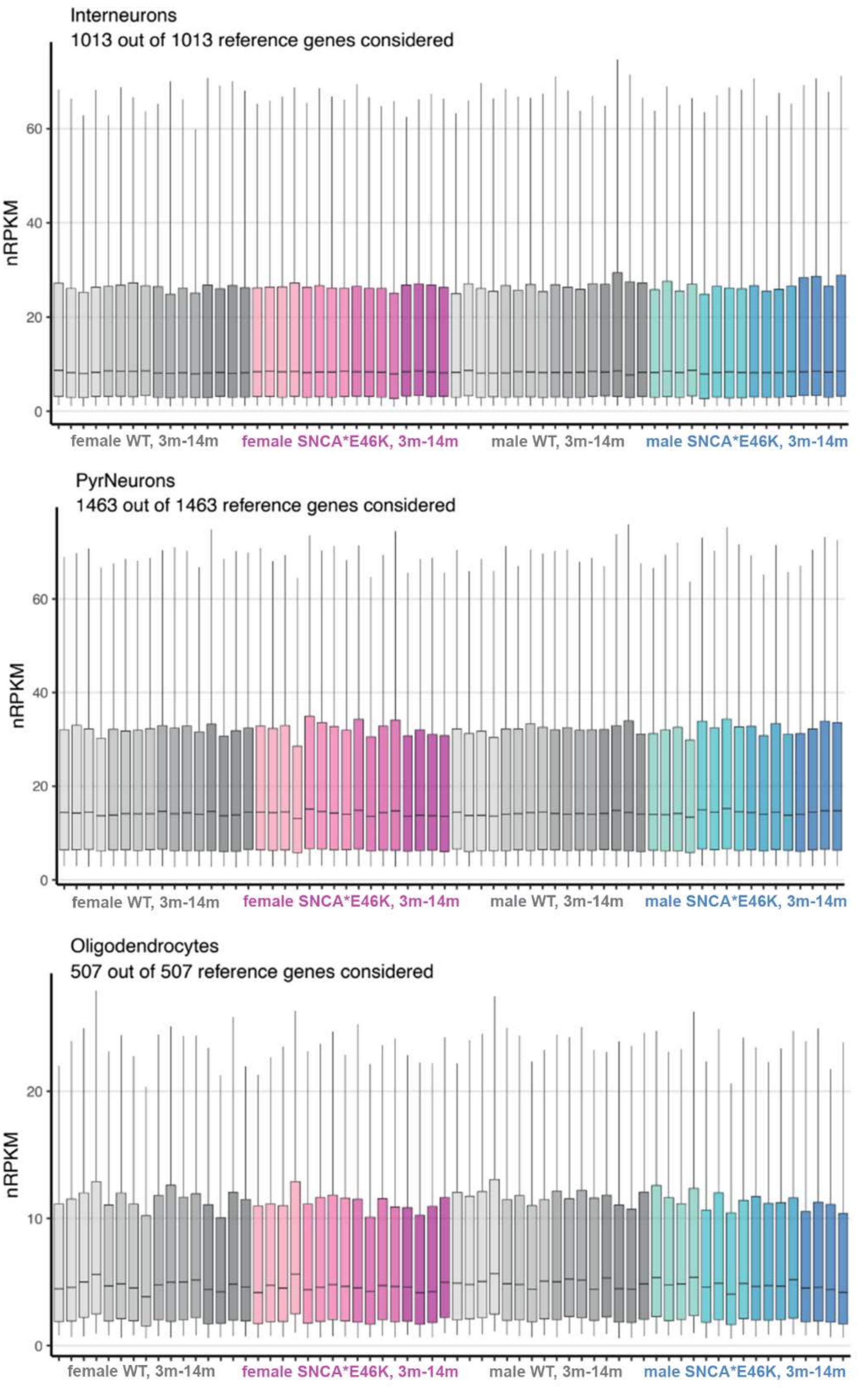
Interneurons
1013 out of 1013 reference genes considered
nRPKM
0
20
40
60
female WT, 3m-14m
female SNCA*E46K, 3m-14m
male WT, 3m-14m
male SNCA*E46K, 3m-14m
PyrNeurons
1463 out of 1463 reference genes considered
nRPKM
0
20
40
60
female WT, 3m-14m
female SNCA*E46K, 3m-14m
male WT, 3m-14m
male SNCA*E46K, 3m-14m
Oligodendrocytes
507 out of 507 reference genes considered
nRPKM
0
10
20
female WT, 3m-14m
female SNCA*E46K, 3m-14m
male WT, 3m-14m
male SNCA*E46K, 3m-14m

24

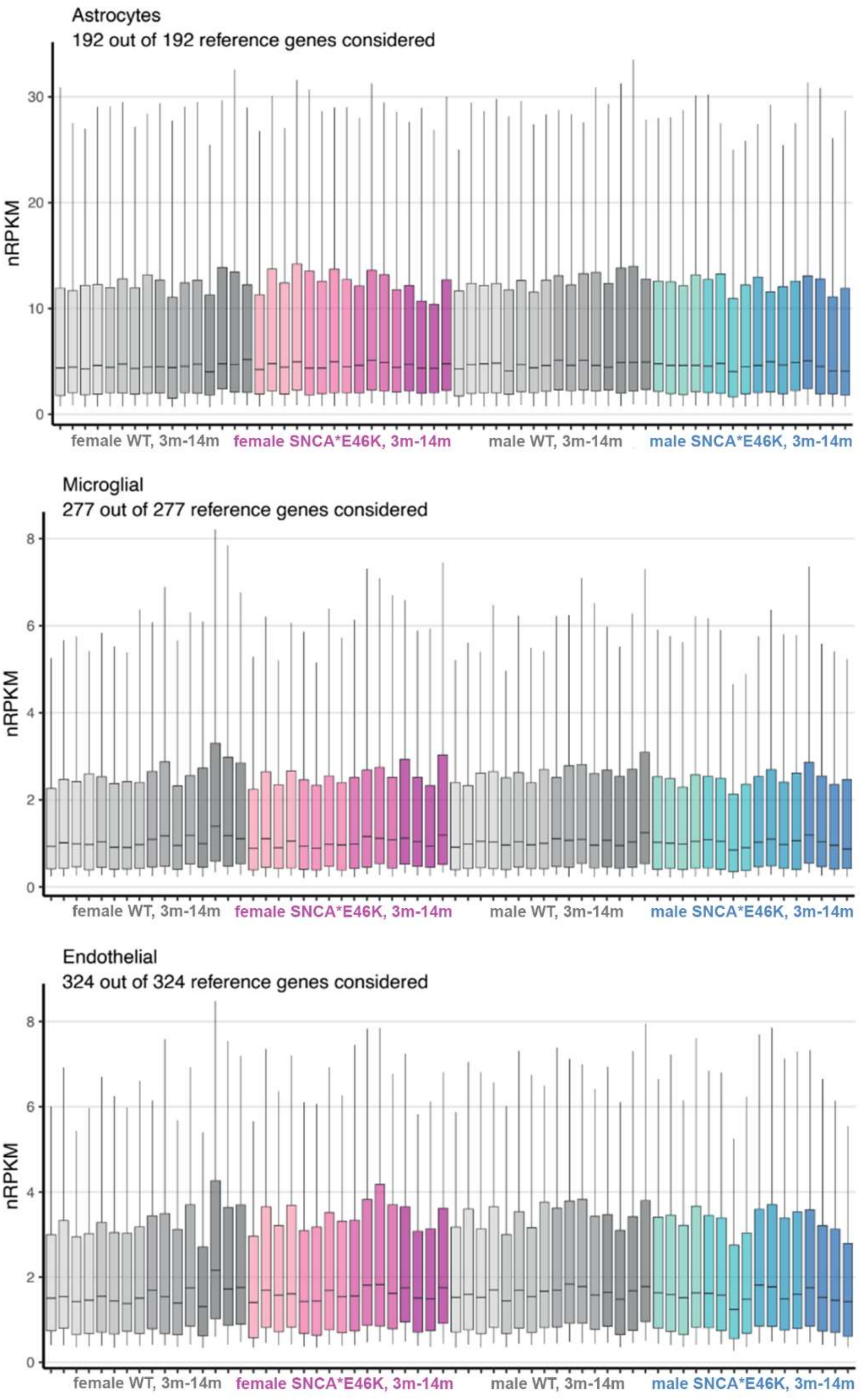


**Suppl. Fig. 4 Cell type-specific expression profiles across all samples as defined by reference profiles**. See main text, Material and Methods section, for details.

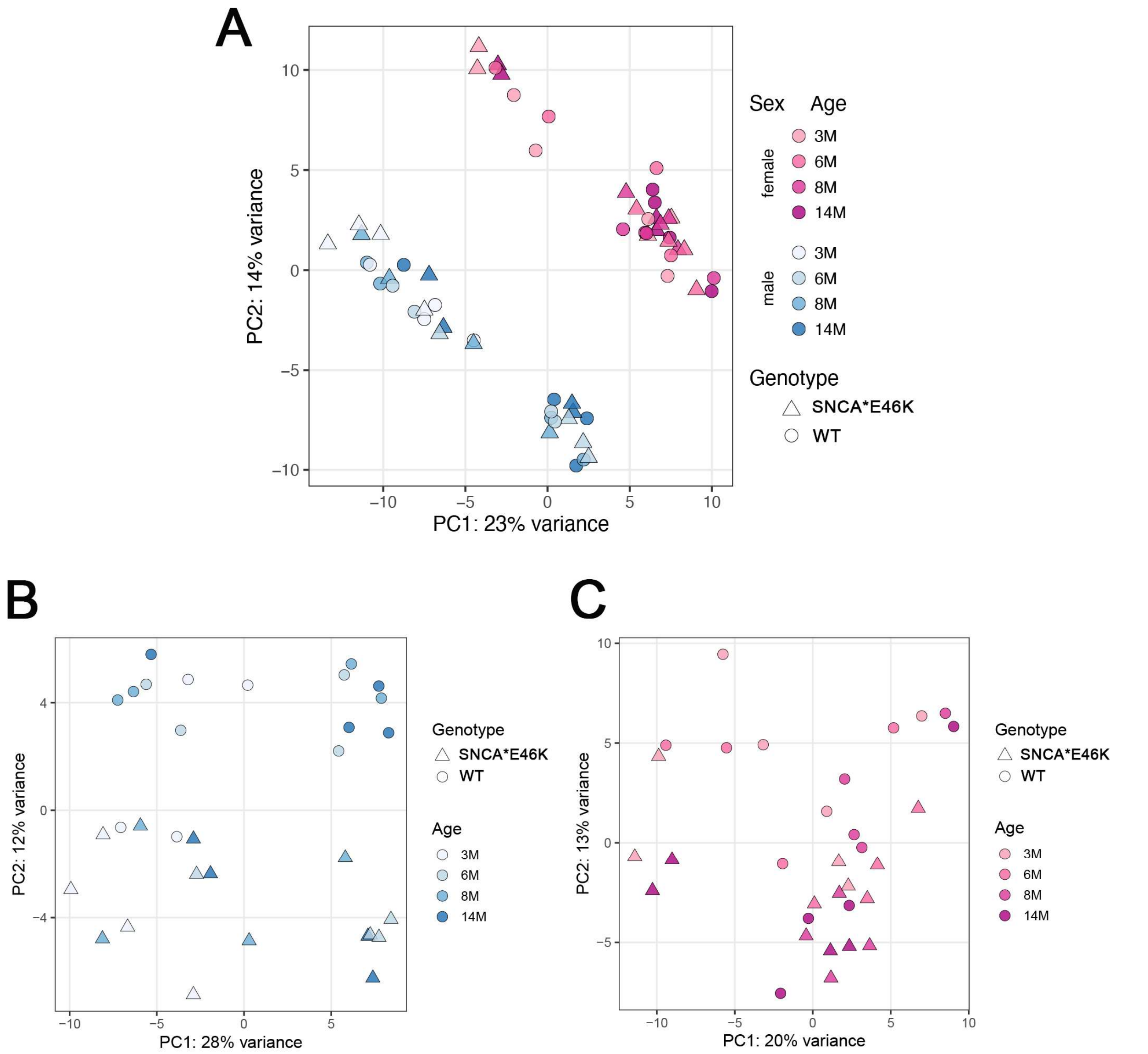


**Suppl. Fig. 5 A)** Dimensionality-reduced representation (Principal component analysis) of hippocampal gene expression profiles for all samples subjected to RNAseq. First and second component of the principal component analysis on the top 500 most variable genes shown. Uncorrected expression data, with no clear separation pattern emerging. **B)** Similar to A, for males only. **C)** Similar to A, for female only. See main text for details.

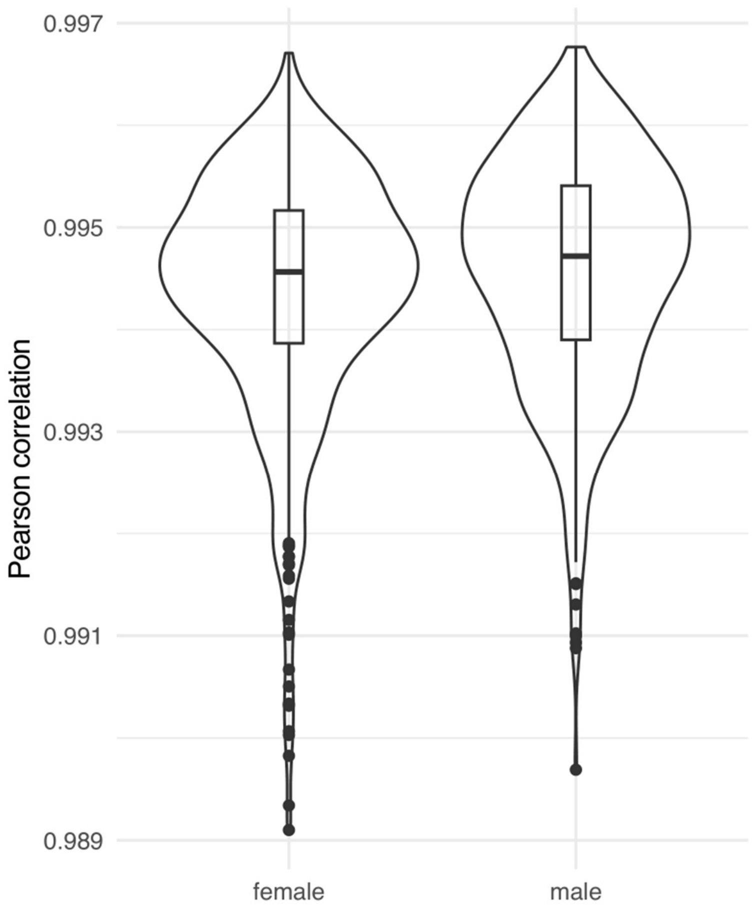


**Suppl. Fig. 6** Violin plots representing the distribution of the correlation coefficients resulting from the pair-wise sample-to-sample correlation for male and female samples. A correlation coefficient close to 1 indicates a positive correlation between samples. The comparable distribution of correlation coefficients in male and female samples indicates that the oestrous cycle is not a relevant factor in hippocampal gene expression.

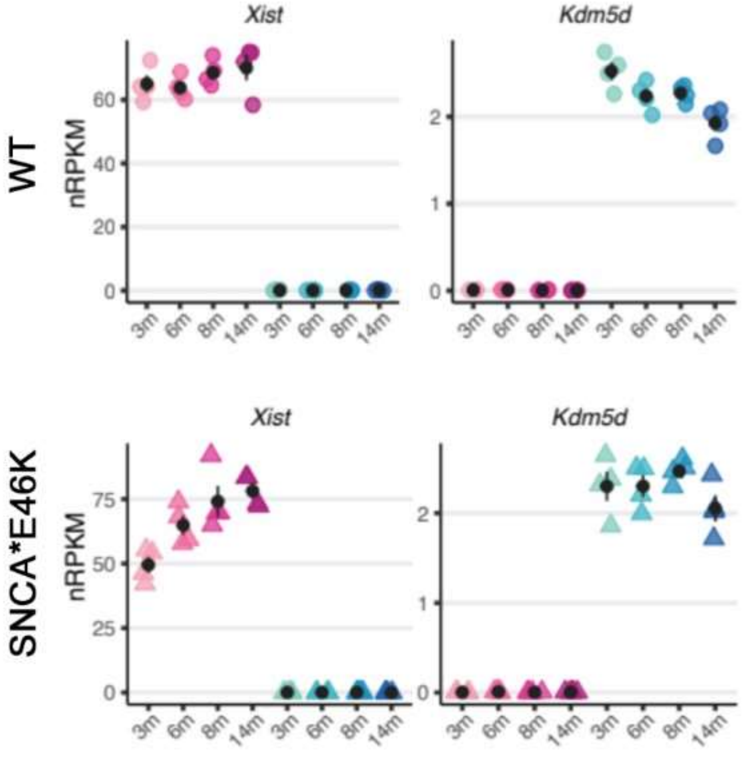


**Suppl. Fig. 7** RNA expression, extracted from RNAseq data, of X-inactive specific transcript (*Xist*¸left) and lysine demethylase 5D (*Kdm5d*, right) in BAC-Tg3(SNCA*E46K) (SNCA*E46K, bottom), mice and wild-type littermates (WT, top). As expected, *Xist* was expressed in females only, while *Kdm5d* in males only, in both genotypes.

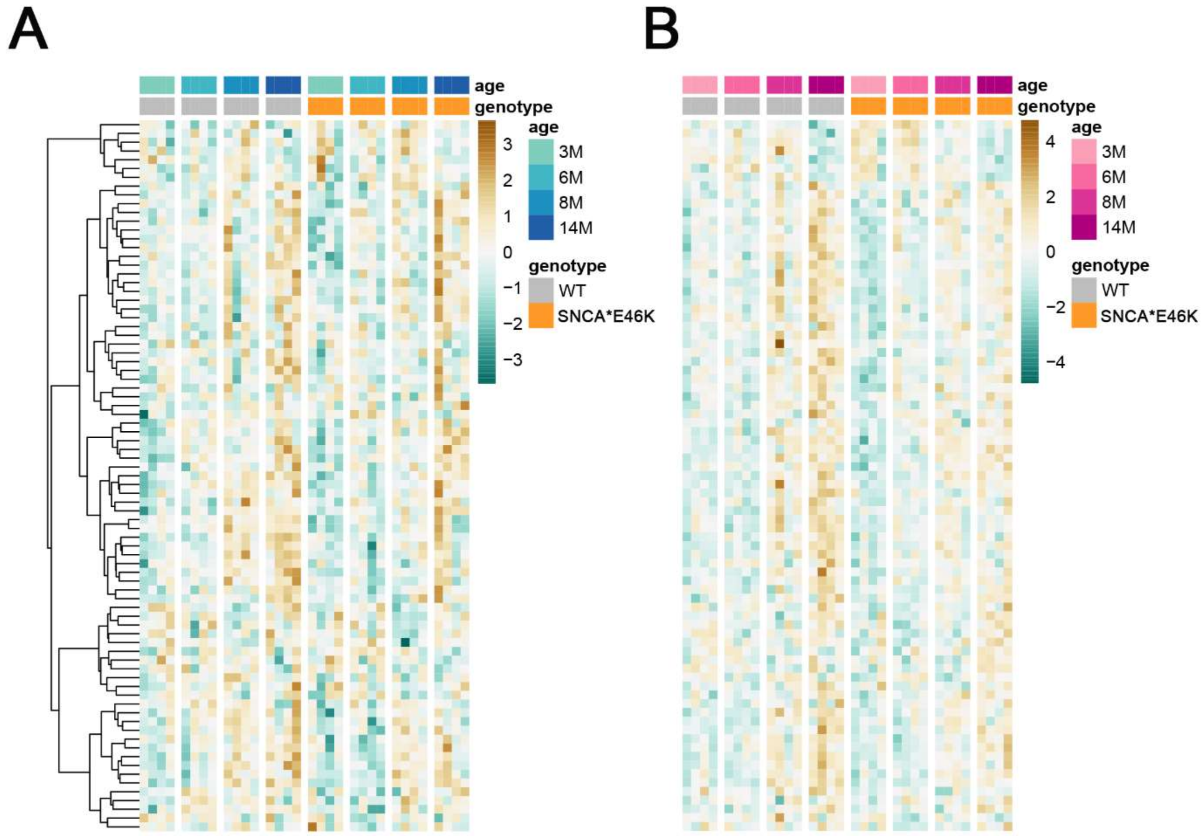


**Suppl. Fig. 8 Hippocampal gene expression pattern in aging WT and BAC-Tg3(SNCA*E46K) mice reflects changes highlighted by the Common Age Signature (CAS)**. **A)** Heatmap showing the expression of CAS genes in male mice. Samples are grouped by age and genotype. **B)** Similar to A, for female mice. See main text for details.
